# Medial Prefrontal Cortex Interacts with the Anteromedial Thalamus in Motivation and Dopaminergic Activity

**DOI:** 10.1101/2020.05.19.104026

**Authors:** Chen Yang, Yuzheng Hu, Aleksandr D. Talishinsky, Christian T. Potter, Leslie A. Ramsey, Andrew J. Kesner, Reuben F. Don, Sue Junn, Aaron Tan, Anne F. Pierce, Celine Nicolas, Coleman B. Calva, Conghui Su, Carlos A. Mejia-Aponte, Dong V. Wang, Hanbing Lu, Yihong Yang, Satoshi Ikemoto

## Abstract

How does the prefrontal cortex (PFC) regulate motivated behavior? Our experiments characterize the circuits of the PFC regulating motivation, reinforcement and dopamine activity, revealing a novel cortico-thalamic loop. The stimulation of medial PFC (mPFC) neurons activated many downstream regions, as shown by functional MRI. Axonal terminal stimulation of mPFC neurons in downstream regions, including the anteromedial thalamic nucleus (AM), reinforced behavior and activated midbrain dopaminergic neurons. The stimulation of AM neurons projecting to the mPFC also reinforced behavior and activated dopamine neurons, and mPFC and AM showed a positive-feedback loop organization. Reciprocal excitatory functional connectivity, as well as co-activation of the two regions, was found in human participants who watched reinforcing videos. Our results suggest that this cortico-thalamic loop regulates motivation, reinforcement and dopaminergic neuron activity.

## INTRODUCTION

The prefrontal cortex (PFC) has been implicated in executive control of goal-directed behavior^1, 2, 3, 4^. Consistently, dysregulation of the PFC is associated with disorders of goal-directed behavior, including drug addiction and depression^5, 6, 7, 8^. Recently, these disorders have been treated with PFC stimulation procedures such as deep brain stimulation (DBS) and transcranial magnetic stimulation (TMS), with promising results^9, 10, 11, 12, 13^. Such treatment effects most likely depend on large-scale brain circuitry regulating the motivation that drives goal-directed behavior^14^, and the details of such circuitry are currently incomplete.

Goal-directed behavior emerges when behavior is reinforced by stimuli/events and can be considered with respect to three key aspects: the motor function (i.e. coordination of movements), the cognitive function (i.e. understanding of the external environment) and the motivation function (i.e. induced arousal that drives attention and action to a goal)^15^. The PFC is thought to regulate these aspects of goal-directed behavior through the basal ganglia^16^. The medial structures of the basal ganglia, including the ventral striatum (VStr) and the ventral tegmental area (VTA), which receive inputs from the medial PFC (mPFC), are thought to be important in the motivational aspect for goal-directed behavior (hereafter referred to as *goal-directed motivation*)^17, 18, 19, 20^. Consistently, mPFC stimulation can activate the VTA-VStr dopaminergic (DA) neurons^17, 21, 22, 23^, whose dysregulation has been implicated in mood and substance-use disorders^24, 25^. Although accumulating evidence strongly supports the claim that the mPFC interacts with the VStr and VTA for goal-directed motivation, the mPFC’s projections to extra-basal ganglia regions may also play a role. While the thalamus has been implicated in goal-directed behavior based primarily on connectivity findings^26^, direct evidence has not yet been provided to support that mPFC projections to the thalamus play a role in goal-directed motivation.

The aim of the present study is to systematically investigate the PFC’s neuroarchitecture in goal-directed motivation. We measured goal-directed motivation using the rate of intracranial self-stimulation behavior (ICSS) in which mice are given the opportunity to respond for optogenetic stimulation of neural populations at cell bodies or terminals. First, we systematically compared PFC areas for ICSS rates, to test which PFC areas are more important in goal-directed motivation^16, 27, 28^, and found that the mPFC supports the highest rates of ICSS among PFC areas. Second, we performed functional MRI (fMRI) with mPFC stimulation, revealing increased activation in various downstream regions including the VStr and extra-basal ganglia regions such as the thalamus and hypothalamus. Third, we found that ICSS was triggered by terminal stimulation of mPFC neurons at various downstream regions, including the anteromedial thalamic nucleus (AM). Fourth, we found that the stimulation of both AM neurons and the AM-to-mPFC pathway supports ICSS and that the AM organizes a positive feedback loop with the mPFC. Fifth, we found that the stimulation of both mPFC-to-AM and AM-to-mPFC neurons activated VTA DA neurons. Finally, we conducted a proof-of-concept experiment providing evidence for the co-activation of mPFC and AM in humans during motivated behavior. In summary, the present study found that the PFC interacts with extra-basal ganglia regions in goal-directed motivation and identified a novel cortico-thalamic positive-feedback loop that can regulate VTA DA neurons and goal-directed motivation.

## RESULTS

### mPFC neurons support high rates of ICSS

The PFC can be divided into three major areas based on connectivity^29, 30^: the mPFC (the infralimbic, prelimbic, cingulate area 2, and medial and ventral orbital regions), orbital PFC (the lateral orbital and anterior insular regions) and motor PFC (the cingulate area 1 and premotor regions). These areas are thought to be important in the motivational, cognitive and motor aspects of goal-directed behavior^16, 31^, respectively. However, it has not yet been documented whether the mPFC is involved in goal-directed motivation more closely than those of the other PFC areas are. To this end, we systematically compared effects of optogenetic ICSS across areas of PFC using the ICSS rate as the measure of goal-directed motivation. In the interest of precision, we distinguished the infralimbic region (IL) from the rest of the mPFC, because previous studies often suggested distinguished role of the IL in motivation and DA activity (e.g.^32, 33, 34, 35^). As an anatomical control, we also examined the tenia tecta (TT), which is positioned immediately ventral to the IL and part of the olfactory system that may be involved in goal-directed motivation^36^.

Each mouse received an injection of adeno-associated viral vector (AAV) with human synapsin promoter (hSyn) carrying channelrhodopsin-2 (ChR2) and enhanced yellow fluorescent protein (EYFP) into one of the areas and had an optic fiber implanted for subsequent photostimulation (Fig. 1a-c). Some mice received the control vector AAV-hSyn-EYFP into the IL. Three weeks after the surgery, each experimentally naïve mouse was placed in an operant conditioning chamber, equipped with two levers (Fig. 1d). Responding on the left “active” lever was rewarded with a train of photostimulation (25-Hz, 8-pulses) in sessions 3-7 (the acquisition phase) and 11-14 (re-acquisition), while no photostimulation was delivered in sessions 1-2 (baseline) or 8-10 (extinction). Responding on the right “inactive” lever produced no consequence throughout sessions 1-14. The mice with IL stimulation markedly increased responding on the active lever in sessions 3-7, decreased lever pressing in sessions 8-10 and reinstated lever pressing in sessions 11-14 (Fig. 1e; Suppl. Fig.1). Response levels on the inactive lever did not increase or change throughout the experiment (data not shown). Control mice, which received IL stimulation, but expressed no ChR2, did not increase lever pressing over baseline levels (Fig. 1e: Control: IL). The data of the control mice established active-lever response levels that determined self-stimulator (>64) from non-self-stimulator, a criterion that is used to interpret ICSS data (mean lever presses plus 3 time their standard deviation; see the method section).

**Fig. 1.**
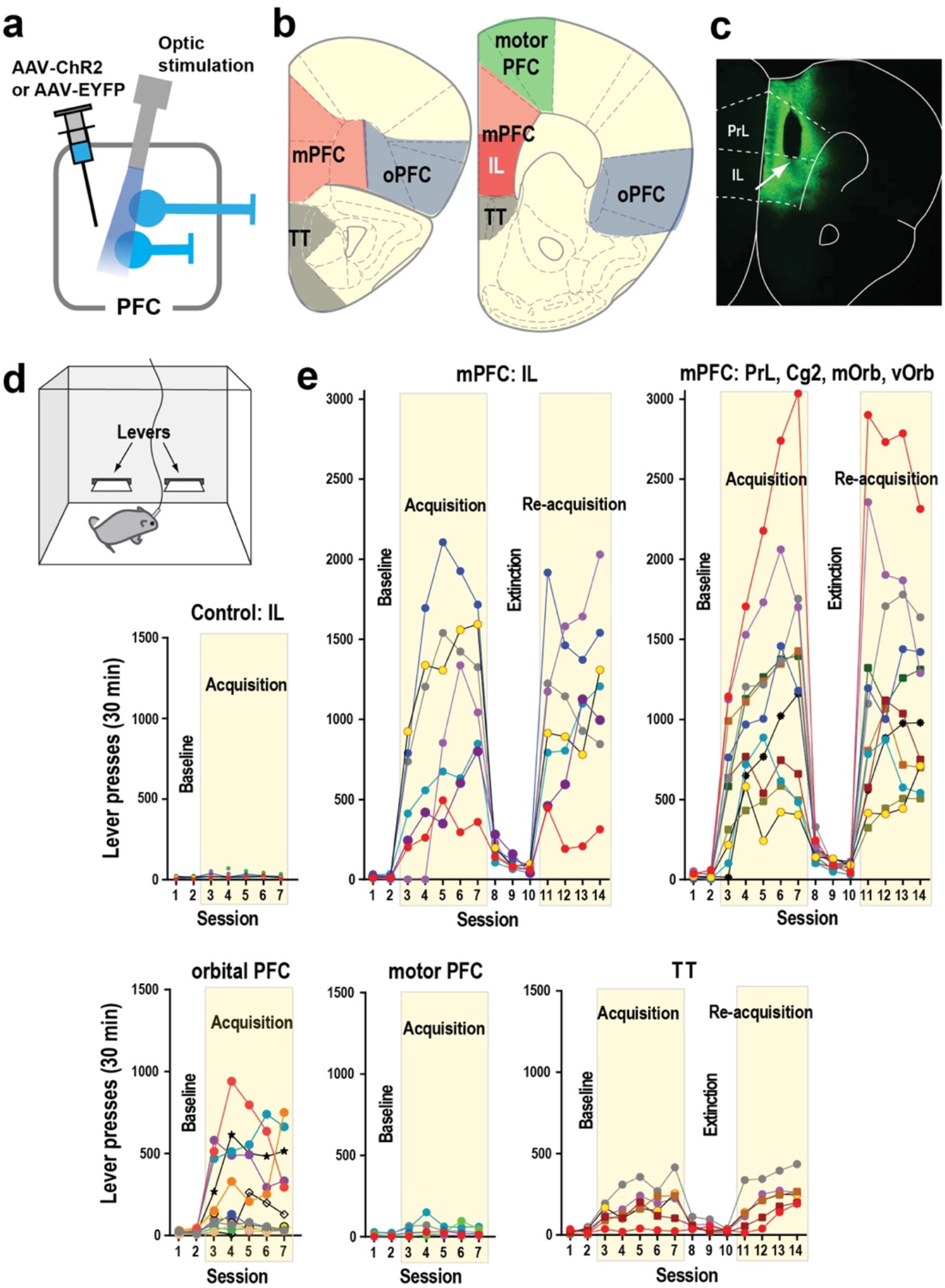
mPFC neurons support high rates of ICSS. **a** C57BL6 mice received an injection of either AAV-hSyn-ChR2-EYFP or AAV-hSyn-EYFP into one of the cortical regions and an optic-fiber implant. **b** Five injection areas. Abbreviations: IL, infralimbic region; mPFC, medial prefrontal cortex; oPFC, orbital PFC; PrL, prelimbic region; TT, tenia tecta. **c** Coronal sections show a representative EYFP expression with the tip of an optic-fiber placement (the arrow) just above the IL. **d** Schematic showing a mouse in a conditioning chamber equipped with two levers for ICSS. **e** ICSS data: Responding on the left lever was rewarded with photostimulation (a 25-Hz 8-pulse train) in sessions 3-7 and 11-14, while no photostimulation in sessions 1-2 or 8-10. Each line shows lever-press data from a single mouse. Abbreviations: mOrb, medial orbital region; Cg2, cingulate area 2; vOrb, ventral orbital region.

Photostimulation at other mPFC regions supported ICSS at rates similar to those of IL stimulation (Fig. 1d; Suppl. Fig.1; a 2^sub-area^ x 5^session^ mixed ANOVA on lever presses: no main area effect, *F*_1,15_ = 0.012, *P* = 0.92). Stimulation of the orbital PFC produced inconsistent effects: It supported ICSS in some mice with moderate rates, while did not in others. By contrast, photostimulation of the motor PFC did not support ICSS. Interestingly, stimulation of the TT reliably supported ICSS, though at low rates. The stimulation of IL supported as vigorous ICSS as that of the rest of the mPFC, and the mPFC as a whole supported ICSS at rates significantly greater than orbital PFC, motor PFC, TT or control group (a 5^group^ x 5^session^ mixed ANOVA on lever presses: a significant main area effect, *F*_4,45_ = 18.48, *P* < 0.0001: *P* < 0.001, a posthoc Tukey test). These results support the idea that the mPFC regulates goal-directed motivation more than other PFC areas. All subsequent experiments focused on the mPFC.

### mPFC-induced motivation is short-lasting and depends on neural firing frequency

The mice with mPFC stimulation performed ICSS with a median response of 1,180 in session 7 (Fig. 1e). That is, half of the mice responded on the lever once every 1.5 s or greater over the course of the 30-min session, and many mPFC mice displayed lever pressing at a constant, fast rate after a few sessions (e.g. Fig. 2a). The highest responder performed at the rate of 1.7 presses every second for a 30-min session, suggesting that mPFC neurons can drive vigorous motivation under optimal conditions (depending on probe placement, ChR2 expression level, etc.). In addition, when reinforcement contingency was reversed between the two levers, the mice quickly learned to reverse response choice from the formerly active lever to currently active lever within a session (Fig. 2b-c), suggesting that the responding was controlled by induced-motivation (i.e. goal-directed emotional arousal) rather than habit formation resulting from repeated fixed-action reinforcement. Moreover, responding became erratic without uninterrupted series of stimulation trains: When two or more responses are required for a single train of mPFC stimulation, which introduces a temporal gap between stimulation, the mice markedly decreased lever-pressing rates, and repeating the procedure did not improve their performance (Fig. 2d). A similar effect was also found with increases in reinforcement intervals, a result described in the fMRI experiment below. Although these observations can be explained by increased physical or cognitive effort, we believe that this is consistent with the idea that stimulation makes the animal seek the next stimulation and that such stimulation-induced motivation is short-lasting. This idea is supported by the next experiment where the effect on ICSS rates diminished with increased photo-pulse interval.

**Fig. 2.**
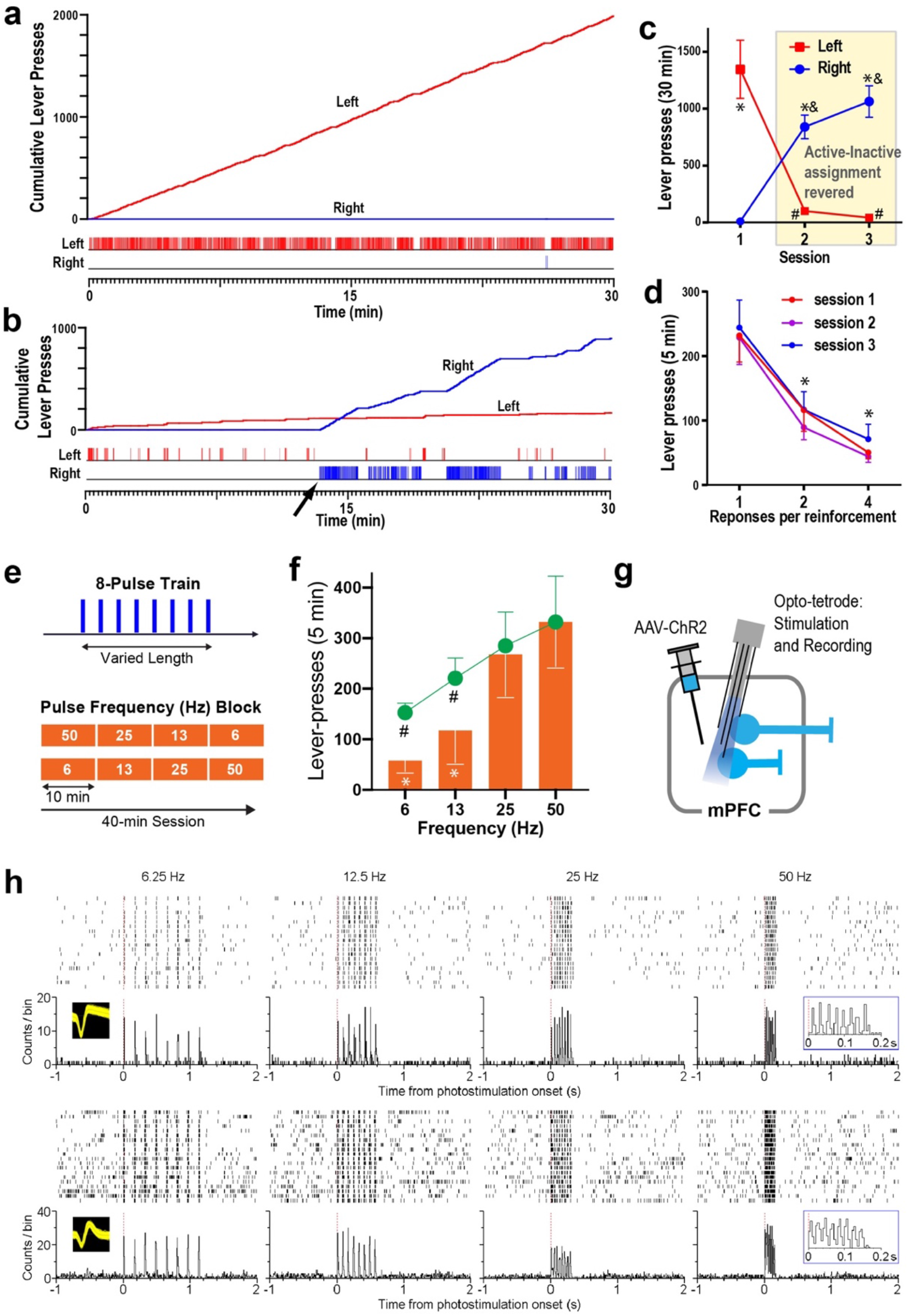
mPFC-induced motivation is short-lasting and depends on neural firing frequency. **a-b** Example of the cumulative record (top) and event record (bottom) showing lever-press responses of a mouse. **b** The cumulative record (top) and event record (bottom) of the mouse (a) that was subjected to reinforcement-contingency reversal where the assignment of active and inactive levers were reversed between the left and right levers for the first time. **c** Mean left and right lever-presses (± SEM; N = 9) before and after reinforcement contingency reversal between the two levers. A 2^lever^ x 3^session^ ANOVA revealed a significant lever-by-session interaction, *F*_2,16_ = 36.9, *P* < 0.0001. * *P* < 0.05, significantly different from the value of the inactive lever-value; # *P* < 0.05, significantly lower than the session-1 value; **&** *P* < 0.01, significantly greater than the session-1 value. **d** Mean lever-presses (± SEM; N = 9) decreased markedly when the mice had to lever-press 2 or 4 times for a mPFC stimulation (a 3ratio x 3 session ANOVA with a main ratio effect, *F*_2,16_ = 19.0, *P* < 0.0001 and no ratio-by-session interaction, *F*_4,32_ = 0.15, *P* = 0.96. * *P* < 0.01, significantly lower than the ratio-1 value with sessions collapsed together. **e** The mPFC was stimulated with 8-pulse trains (top). ICSS was examined over two 40-min sessions where frequencies (6, 13, 25 and 50 Hz) of the train were changed every 10 min in an ascending and descending order (bottom). **f** Active lever-press counts occurring during the second 5-min period of each 10-min block were analyzed with a 2^order^ x 4^frequency^ within-subjects ANOVA. The main frequency effect was highly significant, *F*_3,24_ = 52.31, *P* < 0.00001. Although the order-by-frequency interaction was significant (*F*_3,24_ = 4.29, *P* < 0.05), the effect was small, and the main order effect was not (*F*_1,8_ = 0.09, *P* > 0.05); therefore, the ascending- and descending-order data are combined and presented irrespective of the order factor (orang bars; means ± SEM). The 50Hz stimulation was not significantly more effective than the 25-Hz stimulation, while the 25-Hz stimulation was significantly more effective than the 12-Hz stimulation, which was more effective than the 6-Hz stimulation (Tukey’s test). \**P* < 0.01, significantly different from the 25- or 50-Hz value. The responses for 6- and 13-Hz trains were 38% and 53% of those that they would have responded if they had responded with the same interval for 50-Hz trains (green line graph). #*P* < 0.005, significantly different from the actual responses (multiple t-tests with Benjamini and Hochberg correction). **g** An opto-tetrode probe combining an optic fiber with four tetrodes was surgically implanted for subsequent mPFC stimulation and recording in freely-moving mice. **h** Activities of a putative glutamatergic neuron (top) and a putative GABAergic interneuron (bottom). A close relationship between stimulation frequencies and firing rates was confirmed: the greater stimulation frequencies, the greater firing rates.

We examined how the rate of ICSS changes as a function of stimulation-train frequency (i.e. photo-pulse interval), to determine how important it is for train pulses to occur in a temporarily close manner for motivation. While keeping physical (i.e. lever-pressing action) and cognitive (i.e. knowledge of lever-stimulation contingency) demands and reward quantity (i.e. the number of pulses: 8) constant, the mice received trains with varied pulse-intervals (Fig. 2e). That is, while the amount of reward delivered per response was constant, the speed at which fragments of reward was delivered varied. We found that the mice had significantly higher rates of ICSS rates o for trains with 25 or 50 Hz than 6 or 13 Hz (Fig. 2f). It was expected for the mice to obtain trains at lower rates for low frequency trains, because low frequency trains take longer to deliver than high frequency trains. When the train length was accounted for, the mice reduced response rates significantly more for 6- or 13-Hz trains than for 50-Hz trains (cf. bar and line graphs of Fig. 2f). Therefore, even though the reward amount and effort were constant, the pulse interval determined ICSS rates, suggesting that motivational effect of each pulse decays rapidly within the order of tens of millisecond (see Suppl. Fig. 2 for theoretical consideration). Using an optotetrode probe for simultaneous stimulation and recording, we confirmed that each train can trigger action potentials consistent with its frequency (Fig. 2g-h), suggesting that the frequency of optogenetic stimulation can regulate the firing rate of mPFC neurons. Therefore, the greater firing rates of mPFC neurons, the greater goal-directed motivation.

We conducted additional experiments to further characterize the contribution of mPFC neurons to motivated behavior. First, the inhibition of mPFC neurons induced by the opsin enhanced halorhodopsin 3.0 (NpHR) did not alter preference for environment paired with laser, motor performance, or exploration in an open field, although effectively reduced the number c-fos positive neurons (Suppl. Fig. 3), suggesting that baseline activity of mPFC neurons does not appear to be involved in regulating affective state. In addition, we examined the contributions of mPFC glutamatergic principal neurons and GABAergic interneurons in ICSS, using transgenic mice with AAV-ChR2-EYFP in a Cre-recombinase-dependent double-inverted open reading frame (DIO) construct. Goal-directed behavior was supported by activation of glutamatergic neurons, but not by GABA neurons (Suppl. Fig. 4).

### mPFC stimulation increases activation in downstream regions, including the basal ganglia and others

We performed an fMRI experiment to identify regions downstream of mPFC that are recruited during stimulation that supported ICSS. Rats were used for this experiment because the rat brain provides blood oxygenation level-dependent (BOLD) signals with greater resolution and because it is more difficult to perform fMRI in smaller mouse brains, which are more prone to imaging artifacts. Rats received a unilateral injection of AAV-hSyn-ChR2-EYFP or AAV-hSyn-EYFP into the IL of the mPFC followed by the implantation of an optic fiber above the injection site (Fig. 3a). Rats quickly learned to self-stimulate with mPFC photostimulation (25-Hz 8-pulse train), and then we examined effects of reinforcement intervals: 0, 1, 2, and 4 s, during which the rats could not obtain the next mPFC stimulation upon pressing the lever. Similar to the mice data (Fig. 2d, f), we found that ICSS rates markedly decreased when responding was not reinforced by mPFC stimulation without interruption, in this case increased reinforcement intervals (Fig. 3b), a result consistent with the notion that stimulation-induced goal-directed motivation diminishes quickly within a second. It should be noted that the rates of the rats’ ICSS were generally lower than those of the mice, although all rats did learn to self-stimulate with mPFC stimulation.

**Fig. 3.**
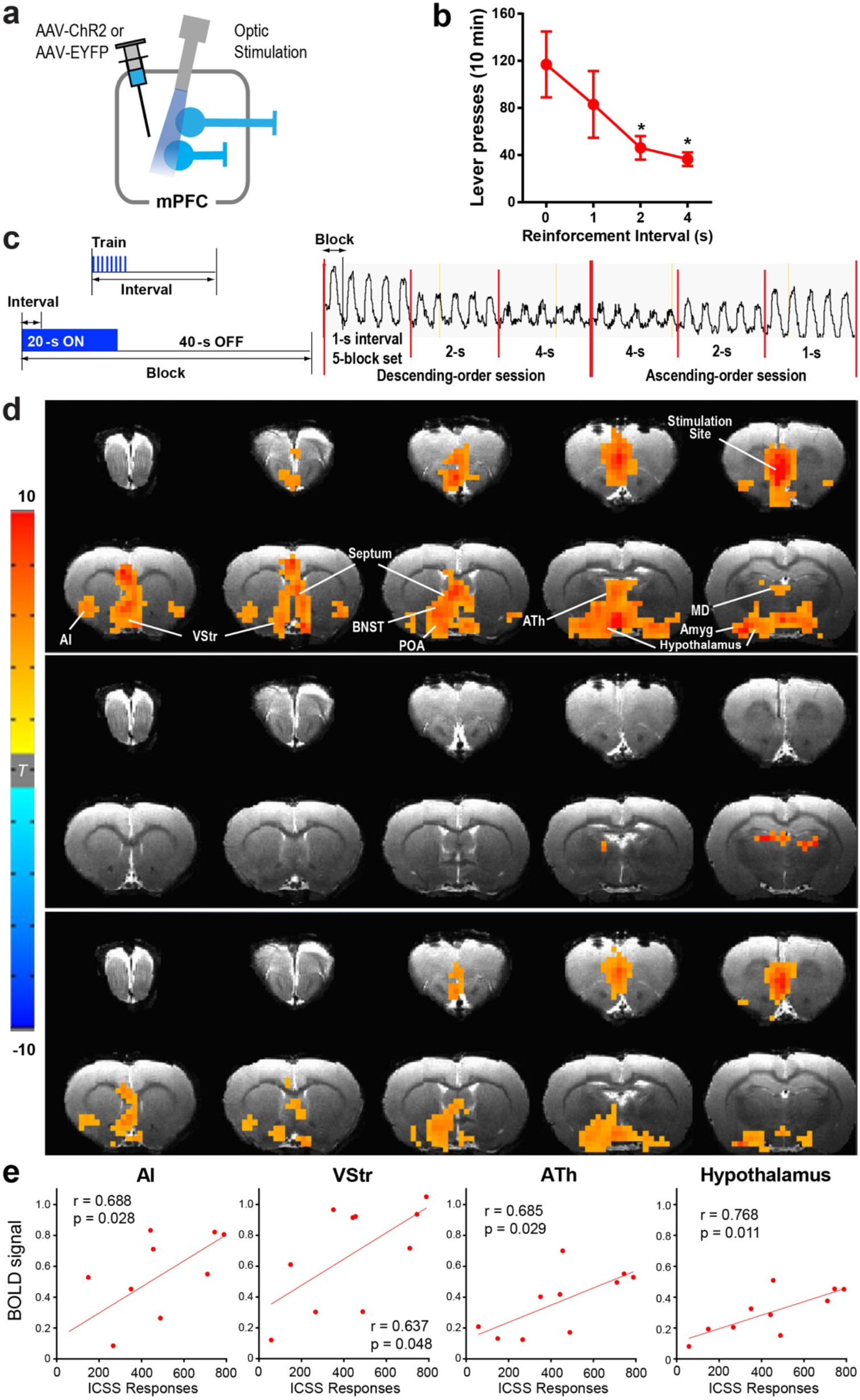
mPFC stimulation increases fMRI BOLD signals in downstream regions. **a** Each rat received AAV-ChR2-EYFP or AAV-EYFP injected into the IL followed by the implantation of an optic fiber at the same region. **b** ICSS: The rats (n = 10) received photostimulation (a train of 8 pulses at 25 Hz) upon lever pressing. After a stable responding over a few sessions, the effects of the reinforcement interval (0, 1, 2, and 4 s) were evaluated with a within-subjects design in that order. Each interval was tested in a 10-min block. Data are mean lever-presses (± SEM). * *P* < 0.05, significantly lower than the continuous-reinforcement schedule (CR) value. **c** fMRI: Photostimulation (a train of 8 pulses at 25 Hz) delivered at the IL with one of three intervals (1, 2 or 4 s) during the 20-s ON phase of a 60-s block. Each block was repeated 5 times for each train interval. Effects of reinforcement intervals were determined in both descending- and ascending-order sessions. Vertical axis indicates raw BOLD signal intensity of one mPFC voxel in arbitrary unit. **d** The unilateral photostimulation with the 1-s interval activated the entire medial network of the PFC, including the medial orbital, prelimbic, ventral anterior cingulate, and infralimbic regions, with stimulation-side dominance. It also elicited significant BOLD signals in extensive subcortical regions of the mPFC (top; n = 10), but limited signals in the control group (middle; n = 7). Significant differences in BOLD signal between the ChR2 and control groups are shown in the bottom panel. Imaging results were corrected for whole brain multiple comparisons (Corrected P < 0.05). Abbreviations: AI, anterior insular region; Amyg, amygdala; ATh, anterior thalamic area; BNST, bed nucleus of stria terminalis; MD, mediodorsal region; POA, preoptic area; VP, ventral pallidum; VStr, ventral striatum. **e** Significant correlations between ICSS responses and the levels of BOLD signals in the AI, VStr, ATh, and hypothalamus. Each dot represents the data of a single rat.

fMRI performed in these rats under anesthesia showed that the interval of unilateral mPFC stimulation determined the strength of BOLD signals (Fig. 3c-d and Suppl. Fig. 5), demonstrating that neural activation elicited by mPFC stimulation decays quickly. Stimulation of the IL increased BOLD signals bilaterally in the entire mPFC, including the IL, prelimbic, medial orbital, and cingulate area 2 regions, an effect consistent with extensive reciprocal connectivity among mPFC neurons within and between the hemispheres^30^ and ICSS data (Fig. 1e; Suppl. Fig. 1). In addition, mPFC stimulation significantly increased BOLD signals in other cortical and subcortical regions including the anterior insular area, tenia tecta, amygdala, VStr, septal area, bed nucleus of stria terminalis, anterior thalamic area, preoptic area, and lateral hypothalamic area. While stimulation-evoked BOLD signals of the mPFC, septum, preoptic area, and amygdala were not significantly correlated with ICSS levels, BOLD signals of the anterior insular area, VStr, anterior thalamus, and hypothalamus were significantly correlated with ICSS levels (Fig. 3e), raising the possibility that these regions integrate goal-directed motivation signals elicited by mPFC stimulation. It should be noted that robust stimulation-evoked BOLD signals found in the visual thalamus and superior colliculus are readily explained by the stimulation of retinal photoreceptors because similar activations were also found in the control group (Fig. 3d; Suppl. Fig. 5) whose retinal photoreceptors must have detected light emitted from the external optical cable due to the testing conducted in complete darkness.

### Axonal terminal stimulation of mPFC neurons supports ICSS in many downstream regions, including the anteromedial thalamic nucleus

Guided by results from our fMRI experiment, we performed ICSS experiments with stimulation in mPFC terminals in downstream regions, to determine the extent to which these pathways are indeed involved in goal-directed motivation. Experimentally naïve C57BL/6J mice received an AAV-hSyn-ChR2 injection into the mPFC and an optic-fiber implantation at one of the downstream regions (Fig. 4a-b). They quickly learned to respond on the active lever that delivered the stimulation of mPFC terminals at sub-cortical regions including the VStr, medial dorsal striatum (mDStr) and internal capsule (IC) (Fig. 4c-d). Because the IC carries cortical information to many downstream regions, ICSS at IC could involve any downstream region. We further dissected the mPFC-IC pathway by placing fibers more precisely in several mPFC terminal fields and found high rates of ICSS with the stimulation of mPFC terminals in the anterior thalamic area (ATh), particularly the anteromedial thalamic nucleus (AM). Of thalamic nuclei, the mediodorsal thalamic nucleus (MD), which is reciprocally linked with the PFC, supported relatively modest ICSS. The reunion thalamic nucleus (Re), which receives mPFC afferents, did not support ICSS. The septum did not support ICSS, either. For hypothalamic stimulation, stimulation sites were placed in the lateral part, since the lateral, but not medial, hypothalamic area is strongly associated with reward ^37^; however, all sites tested in the hypothalamus supported relatively moderate ICSS.

**Fig. 4.**
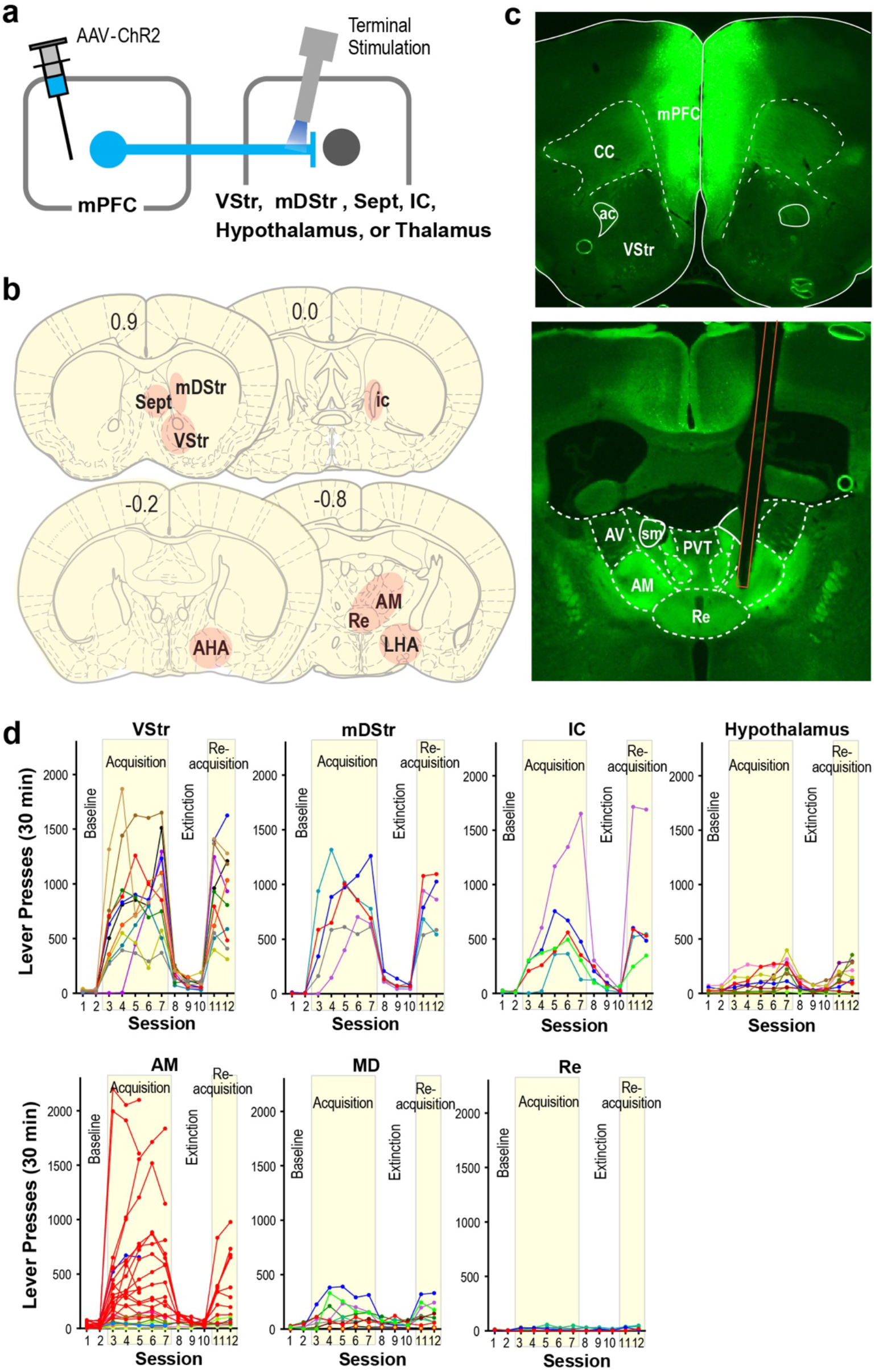
Axonal terminal stimulation of mPFC neurons supports ICSS in many downstream regions, including the anteromedial thalamic nucleus. **a** Diagram showing the site of the injection of AAV-hSyn-ChR2-EYFP and regions of optic fiber implantations. **b** Photomicrograms showing example expression of AAV-ChR2 in the mPFC (top) and in the anterior thalamus (bottom). Orange lines indicate the placement of optic fiber. **c** The sites (n = 80) and effectiveness of axonal terminal stimulation of mPFC neurons. Each dark-gray square indicates the tip of optic fiber, which is accompanied with a colored square below, indicating one of four levels of ICSS rates. Coronal sections were adopted from the mouse brain atlas^68^. Abbreviations: aca, anterior commissure; AHA, anterior hypothalamic area; AM, anteromedial thalamic nucleus; AV, anteroventral thalamic nucleus; DB, diagonal band of Broca; DStr, dorsal striatum; IAM, interanteromedial thalamic nucleus; ic, internal capsule; LHA, lateral hypothalamic area; LS, lateral septum; MD, mediodorsal thalamic nucleus; MS, medial septum; NAc, nucleus accumbens; PC, paracentral thalamic nucleus; PV, paraventricular thalamic nucleus; Re, nucleus of reunion; Rt, reticular nucleus; ST, nucleus of stria terminalis; Sub, submedius thalamic nucleus; VA, ventral anterior thalamic nucleus; VL, ventrolateral thalamic nucleus; VM, ventromedial thalamic nucleus; VP, ventral pallidum; VPM, ventral posteromedial thalamic nucleus. **d** ICSS data of notable regions. An active lever-press was rewarded with photostimulation (a 25-Hz 8-pulse train) in sessions 3-7 and 11-12. Each line shows lever-press data from a single mouse. The data of the AM stimulation site are shown in red, while the data of stimulation sites just adjacent to the AM are shown in different color. Some mice with AM stimulation were only tested for the acquisition phase.

### mPFC neurons projecting to the AM have collateral projections to the VStr and VTA

It is unknown whether and how the AM is involved in motivated behavior. One possibility is that the stimulation of mPFC terminals in the AM activates the collaterals of the same mPFC neurons projecting to the VTA or VStr; therefore, AM neurons may not be involved in the ICSS effects per se. To shed light on this issue, we first examined possible collaterals using an AAV-FLEX-mGFP-2A-SYP-mRuby, which fills the entire Cre-containing cells with mGFP and their terminal boutons with mRuby. We injected this vector into the mPFC and AAV-retro-mCherry-Cre into the AM in C57BL/6J mice (n = 3; Fig. 5a). This procedure should co-label the terminal boutons of mPFC neurons projecting to the VTA or VStr with mGFP and the terminals with mRuby, if mPFC neurons projecting to the AM have collaterals to these regions. One of the mice had its injection located in the anterior thalamic region with little diffusion to the MD or Re (Fig. 5b) and resulted in the cell bodies of mPFC neurons co-labeled with mCherry and mGFP (Fig. 5c). Although the thalamic injection affected not only the AM, but also its vicinity including the anterodorsal, anteroventral, ventral anterior and reticular thalamic nuclei, these structures do not receive afferents from the mPFC^38, 39^. Therefore, mPFC neurons labeled with mRuby are attributed to retrograde activity from the AM. We did find numerous terminal boutons in the VStr and VTA (Fig. 5d-e’), suggesting that mPFC neurons projecting to the AM have collateral projections to the VStr, VTA or both and implying these mPFC neurons in coordinating the activities of these regions.

**Fig. 5.**
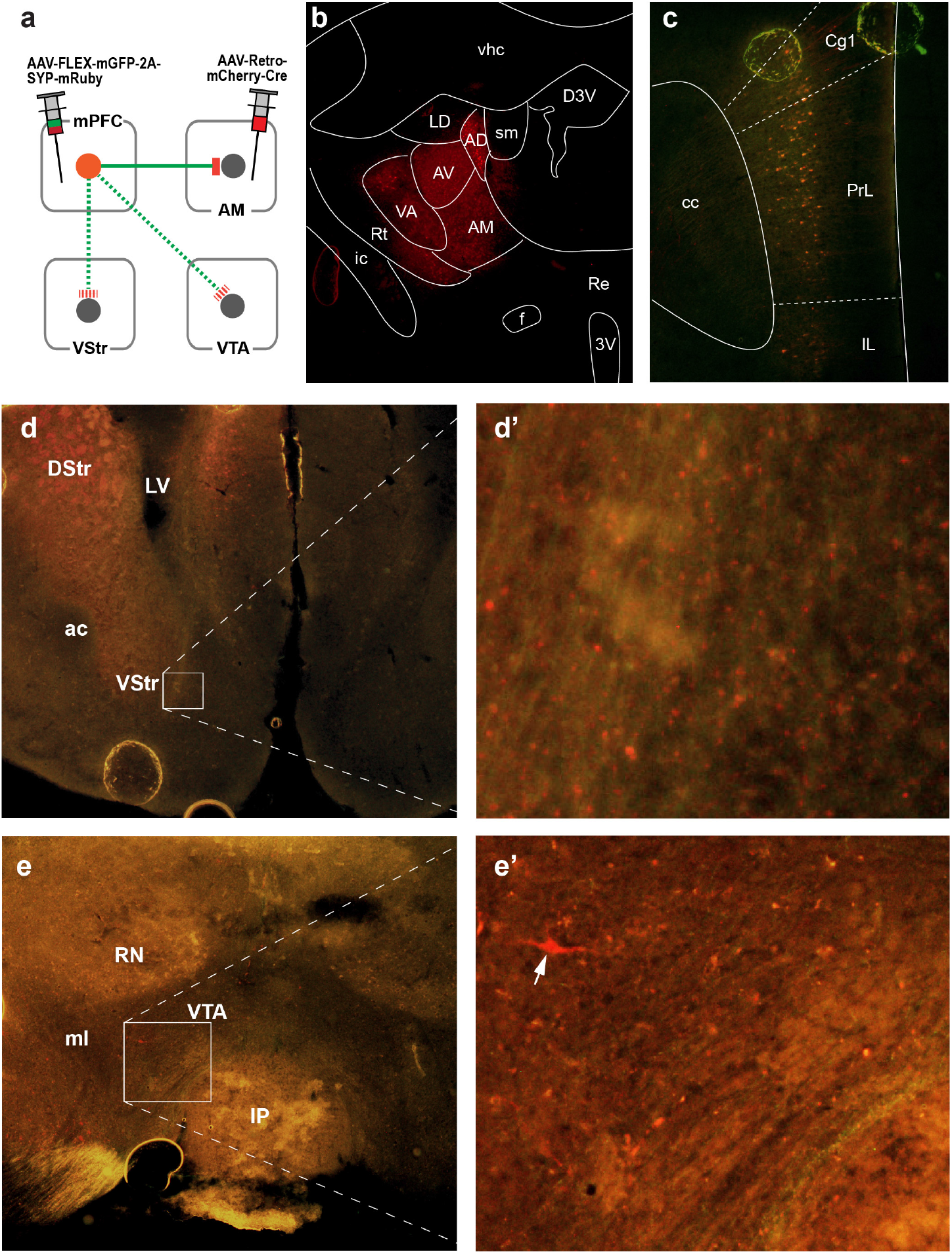
mPFC neurons projecting to the AM have collateral projections to the VStr and VTA. **a** Diagram showing injection manipulations. **b** Photomicrogram showing ATh area affected by injections. **c** Photomicrogram showing double labeled neurons by AAV-retro mCherry-Cre and AAV-FLEX-mGFP-2A-SYP-mRuby in the mFPC. **d - e’** Photomicrograms showing terminal boutons double-labeled by mGFP and mRuby in the VStr (d, d’) and the VTA (e, e’). The arrow indicates a labeled cell without mGFP, suggesting a retrogradely-label by the ATh injection, thereby an ATh-projecting neuron.

### Stimulation of AM neurons supports ICSS

We next examined whether or not direct stimulation of AM neurons supports ICSS. Experimentally naïve C57BL/6J mice received an AAV-hSyn-ChR2 injection into the AM and an optic-fiber implantation at the AM (n = 5; Fig. 6a; Suppl. Fig. 6). They quickly learned to respond on the active lever that stimulated AM neurons. Because AM neurons primarily project back to the mPFC^39, 40, 41^, we then examined whether stimulation of AM terminals in the mPFC supports ICSS. Experimentally naïve C57BL/6J mice received an AAV-hSyn-ChR2 injected into the AM and an optic fiber implanted in the mPFC (n = 5; Fig. 6b). Mice quickly learn to respond on the active lever that delivered photostimulation, suggesting that the stimulation of the AM-to-mPFC pathway supports ICSS. We also examine the same question with an AAV-retro-ChR2 injected into the mPFC and an optic fiber implanted in the AM (n = 5; Suppl. Fig. 6c). The mice did respond for the stimulation, and this result is consistent with the above experiment. However, they decreased responding over the course of the experiment (a significant difference between the first two and the last two sessions), raising the possibility that the AAV-retro-ChR2 had a toxic effect on infected neurons. Overall, these results suggest that AM neurons and AM-to-mPFC neurons support ICSS.

**Fig. 6.**
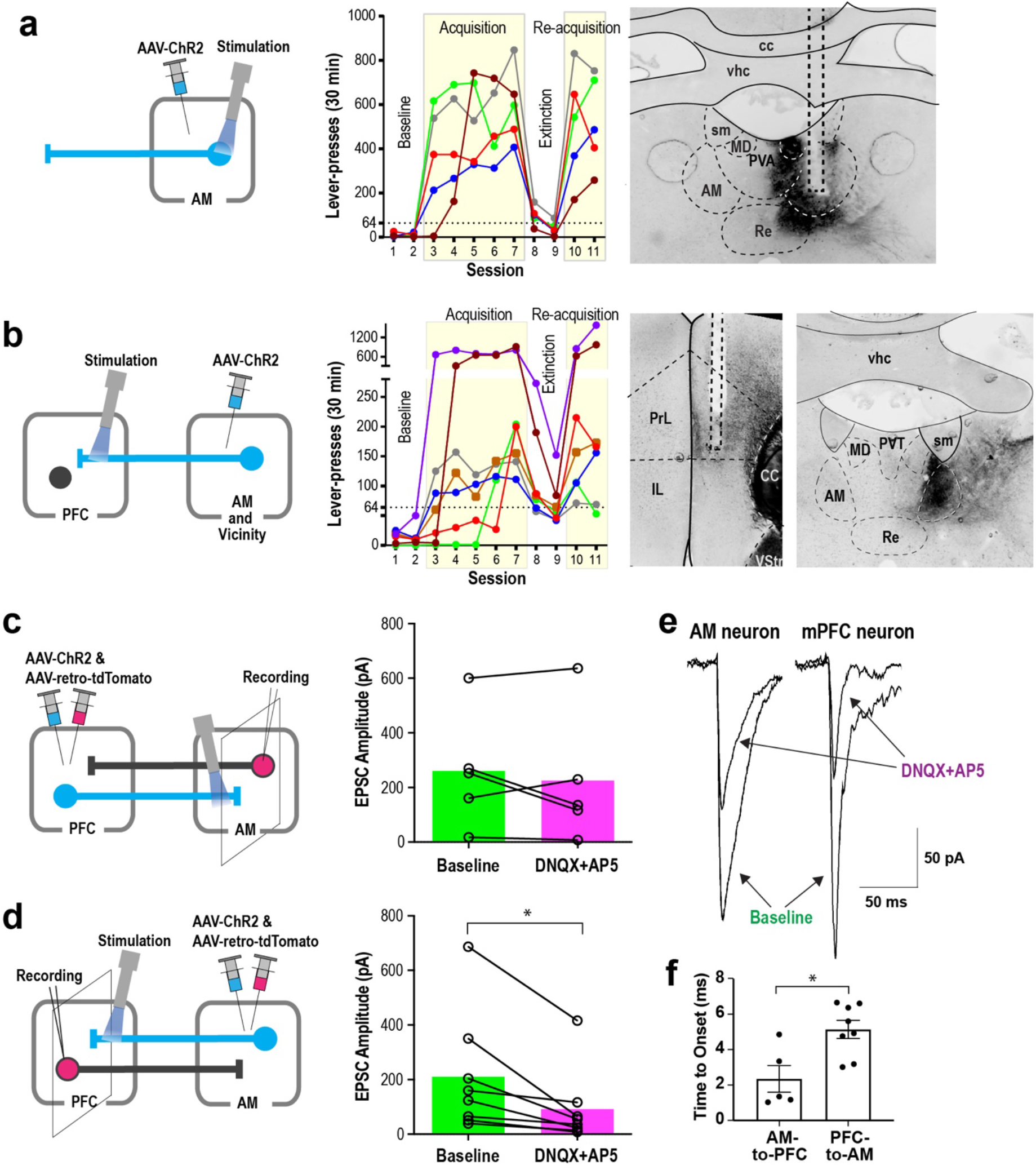
AM neurons support ICSS and form a positive feedback loop with mPFC neurons. **a-b** Left: Diagrams showing the sites of the AAV-ChR2-EYFP injections and optic-fiber implantations. Middle: ICSS data: Responding on the active lever was rewarded with photostimulation (a 25-Hz 8-pulse train) in sessions 3-7 and 10-11, while no photostimulation in sessions 1-2 or 8-9. Each line shows lever-press data from a single mouse. Right: Photomicrograms showing example EYFP expressions and optic-fiber tracks. Occasionally, attenuated expression just below the optic-fiber was found, possibly by photobleaching (a). AM projections were found throughout the mPFC with sparse expression. Stronger anterograde expression was tended to be found in the PrL than the IL and the anterior PrL than posterior PrL, and the two highest responders had their placements in the anterior PrL (b). ICSS with the retrograde procedure supported relatively weak ICSS with the trend of decreased responding over the course of the experiment, especially in the re-acquisition phase; this may suggest toxic effect of the AAV-retro-ChR2. **c,d** Left: Diagrams showing slice electrophysiology preparations with injection sites and stimulation sites. Right: Stimulation-evoked EPSC. * P < 0.05, paired-t test. **e** Example traces: On the left, an AM neuron showing a light-evoked response when adjacent PFC terminals are activated. Response is diminished with application of glutamatergic antagonists. On the right, a PFC neuron showing a light-evoked response when adjacent AM terminals are activated. Response is diminished with application of glutamatergic antagonists. **f** Latencies to response onset as defined as time to achieve 10% of the peak response. * P < 0.01, unpaired-t test.

### AM neurons form a positive feedback loop with mPFC neurons

Many known pathways involved in goal-directed motivation have reciprocal connectivity and function via positive-feedback to promote motivated behavior^18^. To provide evidence for a positive-feedback organization between the mPFC and the AM, we conducted brain slice electrophysiology experiments. Specifically, we examined whether the stimulation of mPFC-to-AM neurons excites AM-to-mPFC neurons and vice versa. C57BL/6J mice received an injection containing AAV-hSyn-ChR2-EYFP and the retrograde tracer AAV-retro-tdTomato into either the mPFC (Fig. 6c) or the AM (Fig. 6d). Of the 11 cells recorded in the AM, 5 were light responsive, confirming that mPFC neurons provide excitatory inputs to AM-to-mPFC neurons (Fig. 6c, e). The application of glutamate receptor antagonists moderately inhibited evoked-EPSC of the AM-to-mPFC pathway in 3 of 5 cells, but not in 2 cells. Of the 8 cells recorded in mPFC, 8 were light responsive, confirming that AM neurons can excite mPFC-to-AM neurons (Fig. 6d-e). Bath application of glutamate receptor antagonists resulted in a 62% inhibition of EPSC of the mPFC-to-AM pathway. Interestingly, EPSC latency to response was faster from AM-to-mPFC than mPFC-to-AM neurons (Fig. 6f; two-tailed t-test, t_11_ = 3.18, *P* < 0.01). Together with the ICSS data, these data suggest a positive-feedback organization between the mPFC and the AM in goal-directed motivation.

### mPFC-to-AM neurons or AM-to-mPFC neurons regulate the activity of VTA DA neurons

VTA DA neurons play a key role in motivation and reinforcement, and the mPFC can regulate VTA DA neurons. Microdialysis studies have shown that electrical stimulation of the mPFC activates the mesolimbic DA system^21, 22, 23^. Recently, optogenetic stimulation of the anterior forebrain projecting to the VTA was shown to excite DA neurons^17^. Because the present study found a role of the mPFC-to-AM projection in ICSS, we used a fiber-photometry calcium-signaling procedure in TH-Cre mice to examine whether the stimulation of the mPFC-to-AM pathway activates VTA DA neurons. Due to its role in motivation and reinforcement^19, 42, 43^, the mPFC-to-VStr pathway was also examined in the same mice (Fig. 7a). We found that the stimulation of both the mPFC-to-AM pathway and the mPFC-to-VStr pathway significantly increased GCaMP signals in the VTA (Fig. 7b; see Suppl. Figs. 7-8 for detailed data and statistical analyses). Notable differences between the pathways were not found, and as long as the optic-fiber placements for mPFC-to-AM stimulation was optimal, mPFC-to-AM stimulation appeared to trigger similar levels of DA neuron activation as VStr stimulation. We then examined whether the stimulation of the AM-to-mPFC pathway activates VTA DA neurons (Fig. 7c) and found that the stimulation of this pathway also significantly increased GCaMP signals in the VTA (Fig. 7d; Suppl. Figs. 7-8).

**Fig. 7.**
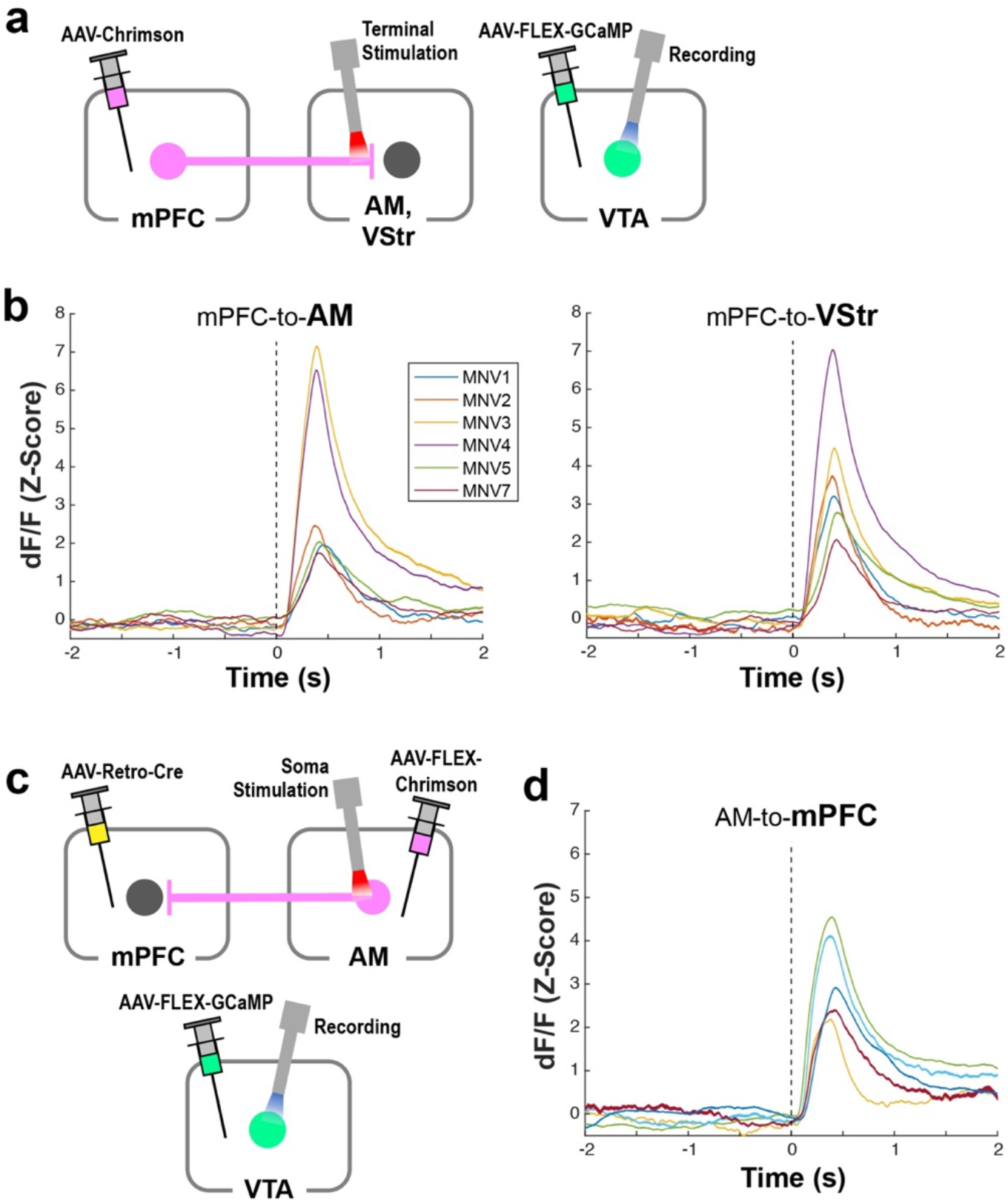
mPFC-to-AM neurons or AM-to-mPFC neurons regulate the activity of VTA DA neurons. **a** Schematic diagram showing AAV-Chrimson injection into the mPFC and stimulation at the AM or VStr (left) and AAV-FLEX-GCaMP injection and recording at the VTA (right) in TH-Cre mice. **b** GCaMP signals (Z-score dF/F) measured in the VTA as a function of 8-pulse photostimulation of the mPFC-to-AM (left) and mPFC-to-VStr (right) pathways in freely moving mice. Colored lines indicate data of 8 mice. The dotted line at 0 s indicates the onset of the photostimulation train. **c** Schematic diagram showing injections of AAV-FLEX-Chrimson, AAV-retro-Cre and AAV-FLEX-GCaMP into the AM, mPFC and VTA, respectively, in TH-Cre mice. Stimulation and recording probes were inserted into the AM and VTA, respectively. **d** GCaMP signals (Z-score dF/F) measured in the VTA as a function of 8-pulse photostimulation of the AM-to-mPFC pathway in freely moving mice. Colored lines indicate data of 5 mice. The dotted line at 0 s indicates the onset of photostimulation train.

### Reinforcing video-clips activate mPFC and AM at the same time in humans

Can our findings have any implication in human motivation? Similar to ICSS in which animals engage in self-initiated responding for stimuli that are not necessary in maintaining biological homeostasis, humans engage in self-initiated, binge responding with stimuli such as substances of abuse and video watching. Recently developed computer-algorithms such as that of the TikTok app can recognize video contents, build a model of what each user likes to watch and show the user more clips according to the model. TikTok-suggested video clips are known to be reinforcing and to induce binge viewing of successive clips^44^. Using fMRI, we examined whether TikTok-clip watching activates the mPFC and AM at the same time in humans. Each human participant watched TikTok video-clips suggested by the TikTok algorithm model developed for the participant as well as TikTok clips assembled randomly without any model.

App-suggested clips activated the AM and BA9 of the mPFC in addition to various other brain regions including both ventral and dorsal visual streams, the frontoparietal network, bilateral arterial insulas, posterior cingulate cortex, parts of middle brain and cerebellum, while they decreased BOLD signals in a set of brain regions including dorsal anterior cingulate cortex, middle cingulate cortex, and cuneus/precuneus, in contrast to rest (Fig. 8a). Control clips also activated similar regions, except the BA9 of the mPFC, with less activation in general (Fig. 8b). The app-suggested clips induced higher activation in regions including the mPFC, AM, temporal gyri and posterior cingulate/precuneus area than the control clips (corrected P < 0.05) (Fig. 8c). The resting-state functional connectivity analysis with AM as seed (Suppl. Fig. 9) shows connections with the entire cingulate cortex area and a large portion of the mPFC (corrected P < 0.05) (Fig. 8d). To determine the region-of-interest (ROI) within the mPFC, we chose the overlapping mPFC sub-area between the zone activated more by the app-suggested clips than the control (Fig 8c) and the AM-connected zone found with the resting-state analysis (Fig 8d), resulting in BA 9 and BA 32 (Fig 8e). Resting-state functional connectivity analysis using the mPFC ROI as seed revealed a significant connection with the AM, in addition to the VStr and VTA, well-known reinforcement/motivation regions, and the PCC and bilateral temporal gyri, which had displayed higher activation with video-clips watching (corrected P < 0.05) (Fig 8f). Further ROI-based psychophysiological interaction (PPI) analysis^45^ showed significantly higher BOLD signal coupling between mPFC and AM upon viewing app-suggested clips, comparing to viewing control ones (PPI effect: mean (SEM) = 0.092 (0.029), P < 0.005). In addition, a dynamic causal modeling (DCM) analysis with parametrical Bayesian model reduction frame^46, 47^ indicates a reliable model (P(m|Y) = 0.70) showing reciprocal excitatory connections between mPFC and AM (Fig. 8g-h). Moreover, the video viewing task modulates the mPFC-AM system primarily through the disinhibition of mPFC (P(m|Y) = 0.67, Fig 8i). These results support the idea that both the AM and the human mPFC (BA9 and BA32) are reciprocally connected and coactivated at the same time during a motivated state.

**Fig. 8.**
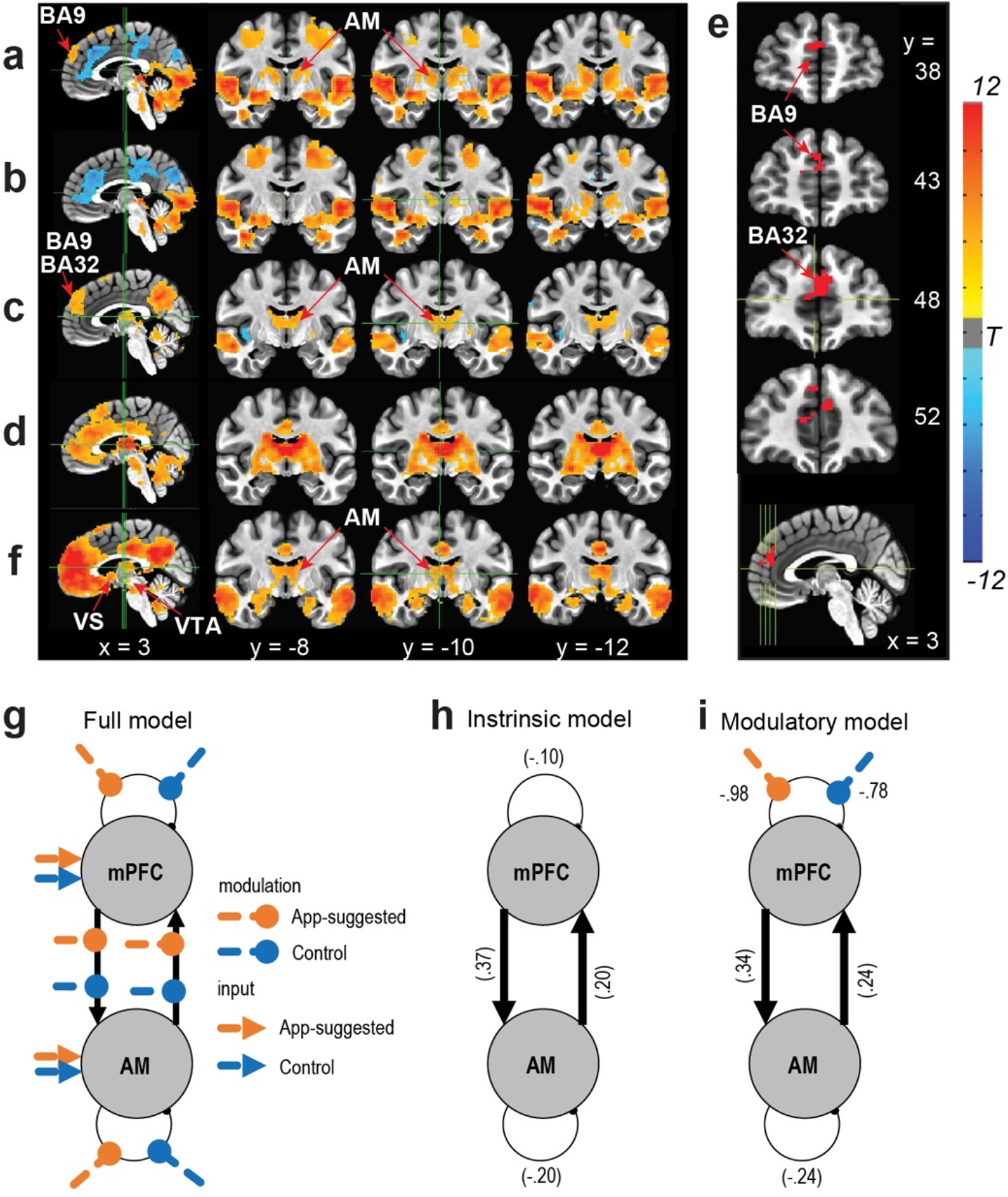
Reinforcing video-clips activate mPFC and ATh at the same time in humans. **a** Change in BOLD signals upon viewing App-suggested video clips. **b** Change in BOLD signals upon viewing control clips. **c** Difference in BOLD signals between app-suggested clips and control clips. **d** Resting-state functional connectivity map with AM as seed. **e** Overlapping mPFC zone (BA 32 and 9) between **c** and **d**. **f** Resting-state functional connectivity map of the mPFC (**e**) as seed. **g** Dynamic causal modelling analysis on the mPFC-AM circuit with a full model structure involving three components consisting of 4 intrinsic connections, 8 experimental modulation on these connections, and 4 direct inputs from experimental conditions. **h** Empirical Bayesian model reduction on the four intrinsic connection parameters indicated the fully connected model was the best intrinsic model (P(m|Y)=0.70) in which mPFC and AM showed excitatory between-region connections and inhibitory within-region connections. **i** Empirical Bayesian model reduction on the sixteen full model parameters showed the best model (P(m|Y)=0.66) in which parameters with posterior probability >0.95 include: 1) intrinsic positive input from mPFC to AM; 2) intrinsic positive input from AM to mPFC; 3) selfinhibition of AM; 4) disinhibition of mPFC by App; and 5) disinhibition of mPFC by controlvideo viewing.

## DISCUSSION

Several novel concepts have emerged from the present study. First, the mPFC regulates motivation for goal-directed behavior via multiple pathways that include extra-basal ganglia regions. Second, the mPFC and the AM interact with each other in a positive-feedback manner for goal-directed motivation. Third, the AM-to-mPFC pathway can regulate DA neuron activity. In addition, the present study provides strong support for the idea that mPFC regulates motivation more effectively than adjacent areas^16, 27, 28^ by providing systematic-comparison data. Before discussing the first three concepts and associated issues, we should discuss behavioral observations on ICSS involving mPFC stimulation, showing how mPFC stimulation is motivating.

Our experiments suggest that mPFC stimulation is highly motivating or reinforcing, and yet this effect dissipates quickly upon the offset of the stimulation. Experimentally naïve mice quickly learned to increase responding on a lever that contingently reinforced by mPFC stimulation. Optogenetic mPFC ICSS rates were essentially as high as those of ICSS with VTA DA neurons that we found in our previous studies using a comparable procedure^48, 49^. This may mean that mPFC stimulation is as motivating or reinforcing as direct stimulation of DA neurons, which are considered a key neuronal population for goal-directed motivation^20^. Consistently, when the reinforcement contingency was switched between the two levers, mice quickly abandoned a lever response that no longer provided mPFC stimulation and switched to a new lever that provided the reinforcement. It is also important to note that, despite vigorous engagement in ICSS with a continuous reinforcement schedule, mice were not able to maintain the rate of responding when they had to make even one extra response for each reinforcement (i.e. mPFC stimulation train) or when they had to wait more than 1 s for the next reinforcement, suggesting the transient nature of mPFC stimulation on motivating effect. The fact that the pulse frequency of the train was also found to be critical in ICSS rates underscores the temporary nature of the stimulation on motivating effect. Such a transient effect of mPFC stimulation on ICSS was parallel to transient effect of BOLD signals induced by mPFC stimulation. Similar transient effects of the stimulation of VTA DA neurons on ICSS have been previously reported^48^.

Our experiments demonstrate that mPFC projections to extra-basal ganglia regions mediate motivation and reinforcement. It is established that mPFC projections to the basal ganglia are important in goal-directed motivation (Fig. 9a): Response-contingent stimulation of the mPFC-to-VStr pathway^19^ or PFC-to-VTA pathway^17^ triggers ICSS, among other findings. Our rodent fMRI procedure revealed that mPFC stimulation increased activation in the VStr. However, the procedure did not detect the activation of VTA, even though robust VTA activation was detected by fiber photometry following mPFC stimulation. We entertain several reasons: First, the MRI receive coil was optimized for the detection of forebrain signals, but not those of inferior brainstem regions including the VTA; second, mPFC stimulation may have activated both VTA DA neurons and GABAergic neurons that inhibits DA neurons, resulting in no BOLD signal; third, the anesthetized rats were used; fourth, interactions of these conditions. Our rodent fMRI procedure also revealed various other downstream regions, including the septum, the thalamus and the hypothalamus. Our optogenetic investigations found that mice engaged in ICSS with the mPFC’s terminals not only in the VStr and dorsal striatum, but also in the internal capsule, which carries cortical signals to subcortical structures, the thalamus and the hypothalamus. The hypothalamus or the mediodorsal thalamic region supported relatively low rates of ICSS, but septal stimulation did not support ICSS. However, we found that the stimulation of the anterior thalamic area, particularly the AM, supported relatively high rates of ICSS.

**Fig. 9.**
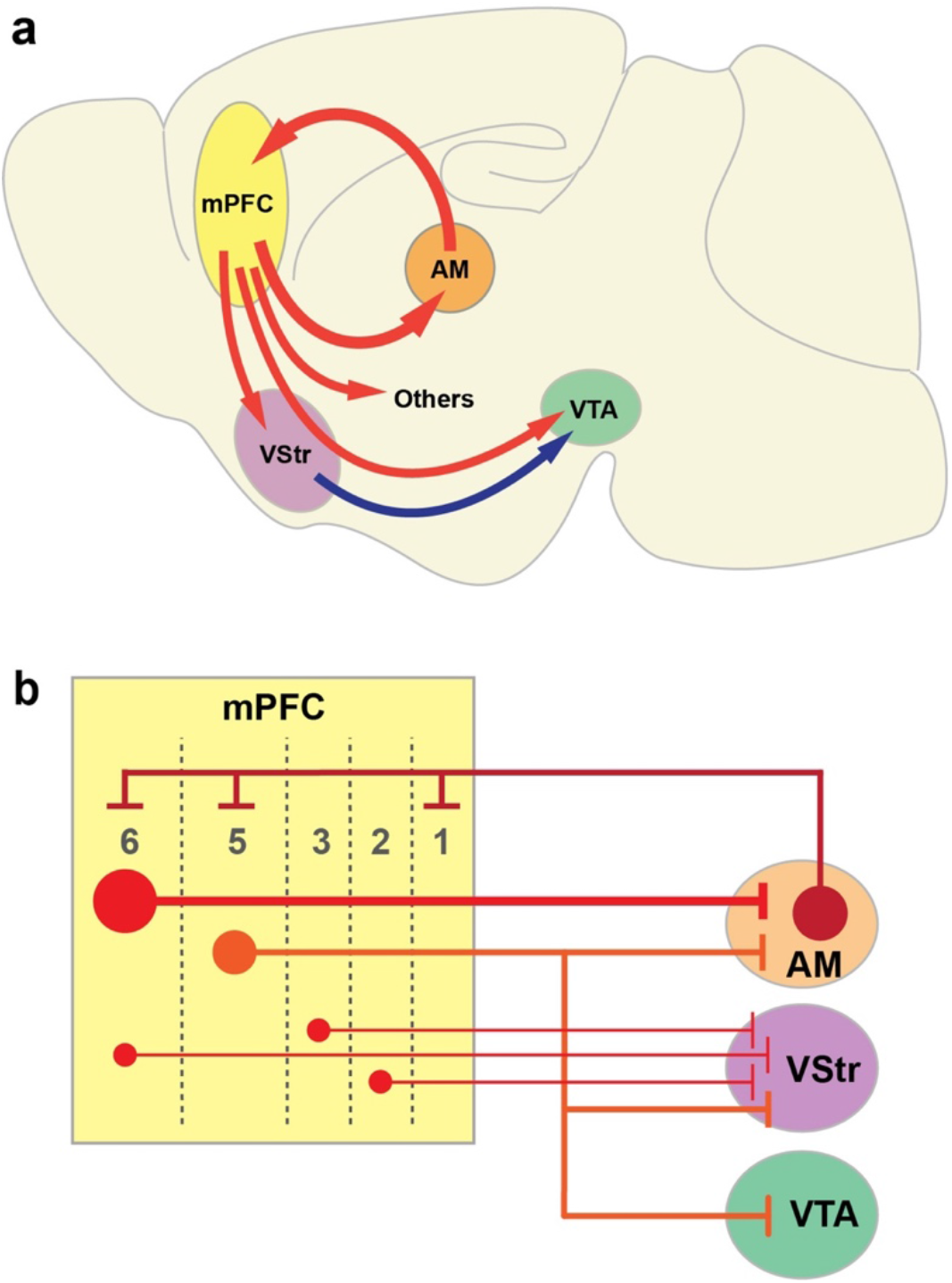
The mPFC regulates goal-directed motivation and dopamine activity via multiple projection pathways. **a** Schematic drawing of sagittal brain section showing the mPFC-AM loop and other key mPFC pathways to regions that participate in regulating goal-directed motivation and DA activity. **b** Schematic drawing showing a microcircuit model between the mPFC and its downstream projection regions of the AM, VStr and VTA.

The present study is the first to demonstrate that the mPFC interacts with the AM in motivation and reinforcement (Fig. 9a). Previous studies implicated the ATh, which includes the AM, in attention^50, 51^. We believe that attention as a descriptive concept of AM functions is consistent with its motivational role, because goal-directed motivated behavior is not tenable without attention (i.e. an essential component of motivation). The present study took the advantage of optogenetics, which can selectively manipulate pathways and thereby enabled us to distinguish the AM from other ATh nuclei because the AM, but not the other ATh nuclei, sends efferents to and receive afferents from the mPFC^38, 40, 41^. We found that response-contingent stimulation of mPFC-to-AM, AM, and AM-to-mPFC neurons triggered ICSS. This positive-feedback arrangement of the mPFC-AM interaction is supported by our slice electrophysiology data and our human fMRI data.

We observed differential activations within the human mPFC as well as interactions among the mPFC, AM, VStr and VTA during the video viewing. This could be explained by functional and structural heterogeneity of the mPFC, whose sub-areas are differentially activated depending on the stimulus that participants perceive, the action that participants engage or both. First, the video clips that the participants watched largely consisted of social stimuli, which may be processed in certain regions of the mPFC^52^. Second, the participants were simply asked to watch videos, but not physically interact with them. Thus, such conditions of the task may have differentially affected mPFC sub-areas as well as downstream regions. In addition, it is important to consider the mPFC layers for understanding how they interact with downstream regions. mPFC neurons have multiple cellular layers, which are differentially connected with the downstream regions (Fig. 9b). The AM receives largest mPFC afferents from layer 6; the VStr receives mPFC afferents from multiple layers; and the VTA exclusively receives mPFC afferents from layer 5^53^. The fMRI task may have selectively activated mPFC neurons in layer 6, which is reciprocally linked with the AM, but has little or no efferents to the VStr or VTA, an idea that could explain the observation that the video watching task activated the mPFC and AM, but not the VStr or the VTA. Although such topic is beyond the scope of the present study, which focused on large-scale circuits, it is important for future research to consider how such micro-circuit organization of the mPFC as well as how the sub-areas of the mPFC regulate motivated behavior in differential circumstances.

VTA DA neurons are known as a key structure involved in goal-directed motivation^20, 54, 55^. Our fiber photometry data suggest that the mPFC-AM loop can regulate DA neurons. We found that the stimulation of not only the mPFC-to-AM pathway, but also the AM-to-mPFC pathway, activated VTA DA neurons. Taken together, mPFC neurons projecting to the AM may activate AM-to-mPFC neurons, and in turn they may activate not only mPFC-to-AM neurons, but also mPFC-to-VTA neurons and even mPFC-to-VStr neurons for the activation of VTA DA neurons (Fig. 9). We propose that the mPFC-AM interaction plays an important role in VTA DA neurons and participates in the global brain network for goal-directed motivation.

Our findings on the AM’s role in motivation corroborate several clinical observations showing that patients with infarcts in or in the vicinity of the AM displayed apathy (also known as abulia), a condition with quantitative reduction in goal-directed behavior^56, 57, 58^. Consistently, other clinical studies suggest that the dysregulation of the mPFC leads to apathy^59^. These clinical observations together with the present observations raise the possibility that the mPFC-AM interaction is a key component of brain network regulating normal goal-directed motivation. It is of interest and potential importance to further investigate how the mPFC-AM loop participates in motivated behaviors and whether the mPFC-AM loop participates in motivational disorders such as addiction and depression, which have begun to be treated with PFC stimulation procedures such as DBS and TMS.

## METHODS

### Animals

Adult male and female C57BL/6J mice were purchased from Jackson Labs (Bar Harbor, ME). Transgenic mice (VgluT1-Cre, VgluT2-Cre, Vgat-Cre) were purchased from Jackson Labs and crossed with C57BL/6J and bred at the NIDA Transgenic Breeding Facility. They were group housed in a colony maintained with a 12h light/dark cycle (lights on at 7 am) and had *ad libitum* access to food and water except during testing. All procedures were approved by the Animal Care and Use Committee of the Intramural Research Program of National Institute of Drug Abuse and were in accordance with the National Research Council *Guide for the Care and Use of Laboratory Animals*. To acclimate animals to experimenters, experimenters handled animals for 5 – 10 min a day for 3 – 4 days with or without fiber cables connected to them prior to the start of behavioral testing.

### Viral vectors

AAVs (Table 1) were obtained from the NIDA Genetic Engineering and Viral Vector Core (GEVVC; Baltimore, MD), Addgene (Watertown, MA) and UNC Vector Core (Chapel Hill, NC).

**Table 1.** List of AAVs

| AAV | Titer (vg/m) | Source |
| --- | --- | --- |
| AAV1-hSyn-ChR2(H134R)-EYFP | $7 \times 10^{12}$ | UNC |
| AAV1-hSyn-eNpHR3.0-EYFP | $1 \times 10^{13}$ | UNC |
| AAV1-hSyn-EYFP | $3.7 \times 10^{12}$ | GEVVC |
| pAAV1-Ef1a-DIO-hChR2(H134R)-EYFP | $7 \times 10^{12}$ | Addgene |
| AAV1-Syn-ChrimsonR-tdTomato-WPRE | $2.3 \times 10^{13}$ | Addgene |
| AAV2-retro-EF1a-DIO-hChR2(H134R)-EYFP | $3.5 \times 10^{12}$ | GEVVC |
| AAV2-retro-CAG-tdTomato | $7.2 \times 10^{12}$ | Addgene |
| AAV1-Syn-FLEX-GCaMP6f-WPRE | $5 \times 10^{12} - 9 \times 10^{12}$ | Addgene |
| pAAV5-Syn-FLEX-rc[ChrimsonR-tdTomato] | $2.0 \times 10^{13}$ | Addgene |
| AAV2-retro-CMV-IE-eGFP-2A-iCre | $4.1 \times 10^{12}$ | GEVVC |
| AAV1-syn1-FLEX-mGFP-2A-SYP-mRuby | $6.9 \times 10^{11}$ | GEVVC |
| AAV2-retro-CMV-IE-mCherry-IRES-Cre | $3.2 \times 10^{12}$ | GEVVC |

### Intracranial surgeries

Mice were anesthetized with either ketamine/zylazine mixture (80/12 mg/kg, i.p.) or isoflurane (1 – 2%) for stereotaxic surgeries. Each mouse typically received unilateral injection of a viral vector (200 – 500 nl) and an optic-fiber implantation unless noted otherwise. The coordinates for viral vector injections are summarized Table 2. The optical fiber was implanted 0.2 mm above the injection site.

**Table 2.** Stereotaxic coordinates for AAV injection sites.

|  | Region | Anterior-Posterior | Medial-Lateral | Dorsal-Ventral |
| --- | --- | --- | --- | --- |
| Cortex | Infralimbic | +1.8 | +0.4 | -3.3 |
|  | Prelimbic | +1.7 – -1.8 | +0.4 | -1.8 – -2.1 |
|  | Medial orbital | +2.3 | +0.4 | -2.7 |
|  | Ventral/Lateral orbital | +2.3 | +0.8 – +1.5 | -2.7 |
|  | Cg1/M2 | +1.8 | +0.3 – +0.9 | -1.0 – -1.4 |
|  | Anterior insular | +1.8 | +2.0 – +2.3 | -2.8 – -3.3 |
|  | Tenia tecta | +2.2 | +0.3 – +0.4 | -3.5 – -4.6 |
| Septum | Septum | +1.1 | +0.3 | -3.5 |
| Striatum | Ventral striatum | +1.1 – +1.3 | +0.5 – +1.2 | -4.2 – -4.9 |
|  | Medial dorsal striatum | +1.1 | +0.9 | -3.5 |
|  | Internal capsule | -0.1 | +1.25 | -3.4 |
| Thalamus | Anteromedial n. | -0.8 | +0.7 | -3.8 |
|  | Mediodorsal n. | -1.1 | +0.3 | -2.7 |
|  | Reunions n. | -0.8 | +0.2 | -4.0 |
| Hypothalamus | Hypothalamus | -0.6 – -2.0 | +0.5 – +0.8 | -4.5 |
| Midbrain | Ventral tegmental area | -3.3 | +0.8 | -4.5 |

### ICSS procedure

Experimentally naïve C57BL/6J mice were individually placed in operant conditioning chambers equipped with two retractable levers, cue lights above the levers, and a house light. They were habituated to the chamber for 30 min with levers retracted, followed by two baseline sessions in which no photostimulation was delivered upon lever pressing. Following these sessions, each mouse was simply placed in the chamber where responding on the “active” lever resulted in intracranial delivery of a 8-pulse light train (473-nm blue-light, 3-ms pulse-duration delivered at 25 Hz, and ~7mW laser power at the tip of stimulation sites), while responding on the “inactive” lever had no programed consequence. A 25-Hz 8-pulse train was used to excite PFC neurons because PFC principal neurons can display spiking activity at 20 Hz or slightly over ^60^. The assignment of active and inactive levers between the two levers typically stayed constant throughout experiments unless stated otherwise. Each session lasted for 30 min, and sessions are typically separated by 1 d.

We used the lever-pressing rates of the control mice to establish non-reinforced response levels. The control mice (n = 8) had the mean of 21.8 presses with the standard deviation (SD) of 13.8 per session during sessions 3-7 (Fig. 1D), which led us to derive 64 (mean + 3 * SD) presses or greater in a 30-min session as ICSS responders.

### ICSS: Reversal of active and inactive lever assignment test

The mice with mPFC implants (n = 9) used in the ICSS experiment described above received a lever reversal test over three sessions in which the assignment of active and inactive levers with respect to the right and left levers was reversed. The lever assignment of the first session was consistent with previous sessions, and the lever assignment was reversed in the 2^nd^ session without any cue. The assignment in the 3^rd^ session was the same as that of the 2^nd^ session.

### ICSS: Frequency test

We examined the effects of stimulation frequency at 6.25, 12.5, 25 and 50 Hz in counterbalance order over a 40-min session with the high responders (>1000 lever presses in 30 min) that were used for the reversal test above (n = 9). Each frequency’s effects were examined in a 10-min period (or 2 bins of 5 min); the number of presses during the second bin was used to represent the effect of the specific frequency. We also calculated how fast each mouse would have responded if they had responded for 6-, 13- and 25-Hz trains with the same interval that they responded for 50-Hz trains: 300-s bin / (response interval for 50-Hz trains + train length of 0.1172, 0.577 and 0.283 s for the 6-, 13- and 25-Hz frequency, respectively).

### ICSS: Lever ratio test

Following the frequency test, effects of response ratio schedules of reinforcement were examined. The mice (n = 9) received mPFC stimulation upon either 1, 2 or 4 lever-presses. The lever-press requirement was changed in an ascending order every 10 min. This procedure was repeated for 3 sessions.

### Electrophysiological recording of MPFC neurons

An optic fiber and 4 or 8 tetrodes were combined to make an optetrode as previously described^61^. C57BL/6J Mice (n = 3) received an injection of AAV-ChR2 into the IL and a movable optetrode just above the IL. Trains of 8 pulses at 6.25,12.5, 25 and 50Hz were delivered at a random order on a variable-interval schedule with a mean of 8.5 s ranged between 7 and 10 s.

### Real-time place-preference test with unilateral photo-manipulation

The same test chamber as described in the above ICSS experiments with levers retracted and was divided into two equal size compartments by placing a Plexiglas barrier with the height of 12 mm from the grid floor. To further help mice to distinguish between the compartments, a 5 kHz tone was continuously delivered when mice were in the right compartment, while a 10 kHz tone was delivered when mice were in the left. Wild-type mice with intra mPFC AAV-NpHR (n = 6) and their control counterparts (n = 6) received continuous photostimulation via the implanted optic fiber while in one of, but not the other, compartments. These mice received photostimulation in the right compartment for the first five sessions, no photostimulation in session 6, and photostimulation in the left for the last five sessions. The assignment of tones to the compartments was kept constant throughout the experiment. Each session lasted for 30 min, and sessions were typically separated by 1 d.

### Effects of the bilateral inhibition of mPFC neurons

Mice (NpHR: n = 5; EYFP: n = 4) were run through a series of behavioral tests to examine the effects of bilateral inhibition of mPFC on various behaviors. For all tests, 3.5 – 5 mW of green (532nm) light was continuously delivered through the implanted optic-fiber for the specified durations. Activity and animal positions were determined automatically using video-tracking software (Noldus Ethovision XT).

#### Real-time place-preference test with bilateral photo-manipulation

The mice were attached to optical lines and placed in a custom-made chamber with a plastic divider separating the chamber into two halves. Visual cues (orange horizontal or blue vertical bars) adorned the sides of the two halves and the orientation of this was counterbalanced. Fresh bedding was used for each session. For the first session (baseline) no light was delivered and for the next three sessions light was delivered when the animal was in the “left” half of the chamber only. Animal position was tracked in real time and time spent on either side of the chamber was automatically analyzed via video tracking software.

#### Forced swim test

A 4000-mL Pyrex beaker was filled with water (25°C) to a depth of 6 inches. The mice were connected to optical lines and gently placed into the water. Green light was delivered for the duration of the 6-min trial. Video was recorded from the side and immobility behavior was automatically scored using video tracking software. Only the last 4 min of the session were analyzed ^62^.

#### Open field test

The mice were attached to optical lines and placed in one corner of a square Plexi-glass arena (42×42 cm with 21×21 cm center) with bright overhead lights. Green light was delivered continuously for the duration of the 5-min session. Video was recorded and the time spent in the center vs the border of the arena was analyzed using video tracking software.

#### Novel environment exposure for c-Fos counting

To test the efficacy of the NpHR mediated inhibition of cells in mPFC using our light parameters, we placed animals for 15 minutes in a novel environment, consisting of a round Plexi-glass bowl shaped container (40 mm in diameter) with clean wood chip bedding and a novel object (small ball-shaped plastic toy). The green light was turned on before animals were connected to the fiber optic lines and was continuous for the duration of the session. We then placed the mice back in their home cage and waited 45 minutes before sacrificing and harvesting brains for c-Fos counting.

#### c-Fos Immunohistochemistry and cell counting

After completion of the behavioral experiments, all mice were intracardially perfused with ice-cold 0.9% saline followed by 4% paraformaldehyde. Brains were coronally sectioned at 40um. Brain sections were processed for immunohistochemistry detecting c-Fos using rabbit anti-c-Fos (1:3000; no. AB152MI, Santa Cruz Biotechnology) and goat anti-rabbit AlexaFluor 594 (1:300; Life Technologies) primary and secondary antibodies, respectively. c-Fos expression was captured with a 5X lens on a fluorescent-microscope/video/computer system and quantified 500 x 500 μm area placed in mPFC at the level of the anterior horns of the corpus callosum. We counted cells in sections corresponding to the tip of the optic fiber and 1 to 2 sections anteriorly before the optic fiber for all mice. c-Fos-ir cells were detected automatically and counted using imageJ (NIH).

### fMRI in rats with mPFC stimulation

Rats were anesthetized with isoflurane and received AAV1-hSyn-ChR2(H134R)-EYFP (n=10) or AAV1-hSyn-EYFP (n=7) injections (0.5 μl) into the infralimbic region of the MPFC and an optic fiber just above the injection site. Stereotaxic coordinates for injections are: AP, 3.2; ML, 0.6; DV, 5.2 mm from the skull surface, and optic fibers were placed 0.4 mm above. Three weeks later, the animals were placed in an operant conditioning chamber and trained to press a lever for optogenetic self-stimulation. A 25-Hz stimulation train consisted of 8 pulses (3-ms pulse-duration) was delivered upon pressing the lever.

The rats then underwent fMRI scanning on a Bruker Biospin 9.4T scanner using a protocol detailed in a previous study^63^. During the scanning, animals were kept anesthetized with a combination of isoflurane (0.5%) and dexmedetomidine hydrochloride (0.015 mg/kg/hr). Block-design optogenetic stimulation was delivered to the right mPFC under three conditions: 25-Hz trains with an interval of 1s, 2s, or 4s, respectively (Fig 3c). Each condition consisted of 5 blocks, and each block consisted of 20s stimulus on and 40s off. Two scan sessions with the stimulus order of A) 1s-2s-4s-interval or B) 4s-2s-1s-interval were performed. The order of scans A and B was counterbalanced between animals. BOLD fMRI data were acquired using a T2*-weighted EPI sequence (TE = 13 ms, TR = 1000 ms, segment = 2, FOV = 35 × 35 mm^2^, matrix size = 64×64, slice thickness = 1 mm, slice number = 15). FMRI data were preprocessed with slice timing correction, head motion correction, spatial smoothing (full-width-at-half-maximum 1.25 mm) and normalization. Voxel-wise whole brain activation was then analyzed using a general linear modelling (GLM) approach, in which the response vector was BOLD signal of each voxel, and predictors were the three boxcar functions (defining the three types of stimulation) convolving with canonical hemodynamic response function (HRF), together with six head motion parameters and low frequency drifts (i.e. linear and quadratic changes with scan time) as nuisance variables.

Independent two sample t-tests were conducted to examine brain areas showing difference in activation between the two groups. Results were corrected for multiple comparisons using randomization and permutation simulation in AFNI’s program 3dttest++ to achieve corrected P < 0.05, which was determined by single voxel p-value <0.005 and minimum cluster size =20 voxels (Fig 3d, Fig S4). Relationship between the fMRI activation and reward behavior was examined using a correlation analysis by correlating beta values of each stimulus type resulting from GLM with lever press measured outside the scanner when animals were awake.

### Human fMRI procedure

#### Subjects

Twenty-five healthy students from Zhejiang University participated in this study (Table 3). All the participants have used the TikTok App and maintained an active account. Most participants (72%) only used the App for video viewing, while the rest used the app for both viewing and publishing their own video clips. Sixty-four percent of the participants had used TikTok for more than half year, and 44% reported that they spent more than one hour on watching videos with this app every day. All the participants had normal or corrected-to-normal vision and reported no neurological diseases. This study was approved by the local ethical committee of Zhejiang University. Written informed consents were obtained from every participant before experiment.

**Table 3:** Demographic information

|  |  |  |  |
| --- | --- | --- | --- |
| Subject number | N = 25: M = 11; F = 14 |  |  |
| Age: mean (std.)/range | 24.1 (2.48) | between 21 and 30 yr old |  |
| Use mode (%) | 72: Viewing only | 28: Viewing & publishing |  |
| Use history (%) | 36: less than 0.5 yr | 32: 0.5 – 2 yr | 32: more than 2 yr |
| Time spend per day (%) | 56: less than 1 hr | 36: 1 – 2 hr | 8: 2 – 4 hr |

#### Procedure concerning video clips

We used the video content-recommendation algorism of TikTok, a popular video app, to examine how individualized content presentation modulates brain activity. We obtained a separate singed consent-form from each participant for privacy protection. We acquired individualized video clips (IVs; 6min long) by singing in the account of each participant and recorded clips (Xiaomi, Model: MI9) right before the fMRI experiment. In addition, we acquired control video clips (CVs; 6 min long), which were obtained through a newly registered account so that the TikTok app had no viewing history information or model for suggestion. The same CVs, which included 29 clips ranging from 5s to 21s were used for all the participants. We confirm that 91% of the participants preferred the IVs over CVs, although the rest preferred CVs over IVs.

#### Experimental procedure during fMRI session

The participants were instructed to be relaxed and watched video clips through an angled mirror and heard the soundtrack through headphones. The presentation of these stimuli was controlled by the software E-prime 3.0 (https://pstnet.com/products/e-prime/). The experiment adopted a block design with IVs and CVs, each type of clips was presented in 6 1-min blocks. The two types of the 6 blocks were presented alternatively and separated by a 30-s break, during which the participants viewed a white fixation on a black background. Half of the participants started with an IV block first, while the other started with a CV block first. To minimize disruption, no additional question or interruption was introduced during the experimental session.

#### MRI data acquisition and preprocessing

MRI data were collected using a Siemens 3.0-T scanner (MAGNETOM Prisma, Siemens Healthcare Erlangen, Germany) with a 20-channel coil. High resolution anatomical images were acquired using a T1-weighted magnetization prepared rapid gradient echo sequence with parameters below: TR = 2300 ms, TE = 2.32 ms, voxel size = 0.90 × 0.90 × 0.90 mm^3^, flip angle = 8°, field of view = 240 mm^2^, voxel matrix = 256 × 256. FMRI data were collected using a T2*-weighted gradient echo planar imaging sequence with multibands acceleration (TR = 1000 ms, TE = 34 ms, slice thickness = 2.50 mm, voxel size = 2.50 × 2.50 × 2.50 mm^3^, voxel matrix = 92 × 92, flip angle = 50 °, field of view = 230 mm^2^, slices number = 52, MB-factor = 4). An 8-minute resting-state fMRI data (480 scans) were collected after structural image acquisition, during which participants were instructed to watch a white fixation on a black screen with relaxation, not to think anything in particular and keep still during the scan session. A total of 1095 scans for the task fMRI data were collected using the same acquisition parameters.

Preprocessing of fMRI data included the following steps. First, slice time correction and head motion correction were performed using AFNI (https://afni.nimh.nih.gov/). Then, tissue segmentation was then conducted to extract brains using SPM12 (https://www.fil.ion.ucl.ac.uk/spm/). Structural and functional images were co-registered and normalized into the MNI space using ANTs (http://stnava.github.io/ANTs/). Finally, spatial smoothing was applied to the normalized fMRI data with a 5 mm full-width-at-half-maximum Gaussian kernel. For resting-state data, two more preprocessing steps were included: 1) nuisance variable regression including six-rigid head motion and their forward derivates, FD (see below), and the first 5 principle components from white matter and cerebral spinal fluid (CSF) separately; and 2) a band-pass filtering (0.01Hz – 0.1Hz) was applied.

#### Head motion handling

Follow a previous study by Power and colleagues^64^, framewise displacement (FD, Eq. 1) of fMRI data was calculated for each participant as indices of head motion for task and resting-state data separately.

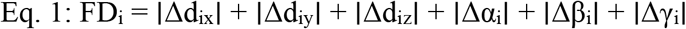

where Δd_ix_ = d_(i-1)x_ − d_ix_, and similarly for the other rigid body parameters [d_ix_ d_iy_ d_iz_ α_i_ β_i_ γ_i_]. A participant would be excluded from statistical analysis if her/his mean FD >0.3mm, or the total number of frames with FD > 0.5mm is more that 10% of the total length of the data. With these two criteria, two subjects were excluded from task fMRI data analysis, and one subject was excluded from resting-state fMRI statistical analysis. Head motions parameters were also included as nuisance variables in the first level fMRI data analysis.

#### First level task fMRI statistical analysis

For the first level analysis, general linear modelling (GLM) was conducted using the command 3dDeconvolve within AFNI. The 6 encoding blocks for IVs and 6 for CVs during encoding were convolved with hemodynamic function to create 2 regressors to assess activity elicited by the two conditions, respectively. Further, two event-related regressors were created to model the transient effect associated with the start and end of video blocks for IVs and CVs, respectively. The six head motion parameters were included in the model as covariates, together with 8 polynomial variables to remove task-unrelated artifacts.

#### Group level task fMRI statistical analysis and results

Voxel-wise one-sample t-tests on beta maps of the block regressors for IVs and CVs were conducted to assess brain activation related to the two conditions, respectively. Paired wise t-test was conducted to examine brain areas showing difference in brain activation under the two conditions. Results were corrected for multiple comparisons (corrected P < 0.05, Fig 8a-c) using randomization and permutation simulation in AFNI (single voxel p-value <0.001 and minimum cluster size =33 voxels).

#### Resting-state functional connectivity of AM and mPFC

Although higher brain activation in anterior thalamus was seen in the paired t-test, the main purpose of the present analysis was to investigate the implication of same mPFC-AM circuit in human motivational behavior. Therefore, we defined AM (Suppl. Fig. 8) by drawing ROI based on the “Atlas of Human Brain” 4^th^ edition^65^. Five voxels in each hemisphere were identified with a spatial resolution of 2.5mm^3^. Time course was extracted from the AM seed and correlated with the time course throughout the whole brain. Resultant correlation coefficients were transferred into Z-maps with Fisher’s Z-transformation (Eq 2). The mPFC seed (Fig. 8e) was defined by the union of AM-seeded restingstate connectivity map (Fig 8d) and task activation region showing higher activity in the IVs condition (Fig 8c). One-sample t-test was apply to the Z-maps and results were corrected for multiple comparisons (corrected P < 0.05, Fig. 8d, f).

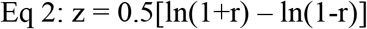

#### PPI analysis of mPFC-AM circuit

Psychophysiological interactions (PPI) is based on statistical models of factorial design by substituting brain activity in mPFC regions for one of the factors^45^. Another factor is the categorical video type condition (IVs vs CVs) in the present study. Equation 3 below illustrates the PPI model.

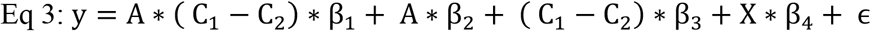

The effect of video viewing condition is assessed with the contrast term (C_1_ − C_2_), where C_1_ and C_2_ were coded as 1 to represent IVs and CVs viewing conditions, respectively. The term A is a physiological variable of mPFC (BOLD signal here), and similarly, y is the BOLD signal of AM regions. While A*(C_1_ − C_2_) indicates the PPI term, its regression coefficient β_1_ is used infer the significance of PPI effect. The term X denotes all nuisance variables and ε is the error term. In the present study, nuisance variables include 6 head motion parameters and 8 polynomial variables determined by 3dDeconvolve, and BOLD signals from the dACC region that showed deactivation during task. Note that PPI and DCM analyses were conducted using SPM12. To keep consistency with results obtained with AFNI, the 8 polynomial variables were included as nuisances and the cut-off for high pass filtering was set to 1024 seconds instead of using the default 128 seconds when running GLM for PPI and DCM.

#### DCM analysis of mPFC-AM circuit

Dynamic causal modelling (DCM) is a technique to infer neural processes that underlie observed time courses^66^, such as fMRI data. The DCM procedure involves the Bayesian estimation of parameters of neuronal system models from which BOLD signals are predicted through converting the modelled neural dynamics into hemodynamic response based on the neurovascular coupling model. Statistically, DCM models the temporal evolution of the neural stats vector as a function of the current state (z), the task input (u) and parameters (*θ^n^*) that define the functional architecture and interactions among brain areas at the neuronal level (Eq 4).

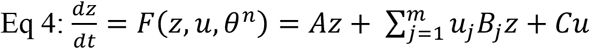

These parameters describe the nature of the three causal components which underlie the modelled neural dynamics: 1) the parameter set A in Eq 5 describes intrinsic effective connectivity among brain regions (a *k* x *k* matrix), 2) the parameter Bj (Eq 6) describes context dependent changes in effectivity induced by the *j*th input uj (a *k* x *k* matrix for each input), and 3) the parameter C (Eq 7) describes direct task inputs into the system that drive regional activity (a *k* x *m* matrix), whereas *k* is the number of nodes in the system and *m* is the number of task inputs.

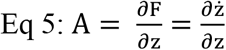

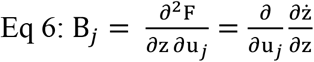

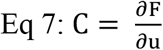

In the present study, we are interested in the mPFC-AM system implicated in motivational behavior. The anatomic definitions for mPFC and AM were the same as we used in the restingstate connectivity analyses (Suppl. Fig. 8), and the task inputs were defined as the two video viewing conditions (i.e. IVs and CVs), yielding a full model structure including 16 parameters (Fig 8. g). We first asked the question of whether human imaging data would produce the same functional architecture revealed with animal physiological techniques by applying the Parametric Empirical Bayesian (PEB) frame^46, 47^ to the intrinsic model (i.e. the field A). The resultant best model indicates reciprocal excitatory connections between mPFC and AM (Fig 8h), which is consistent with the results from animal data showing positive feed-back in the mPFC-AM loop (Fig. 6). Then we asked the second question of how the video viewing condition modulates the system by applying PEB to the full model parameters (i.e. the fields A, B and C). The resultant best model indicates that the task condition modulates the system primarily by dis-inhibiting mPFC (Fig 8i).

### Slice electrophysiology procedure

C57BL/6J mice received injections of AAV-hSyn-ChR2-EYFP and AAV-retro-CAG-tdTomato into either the mPFC (n = 4) or the AM (n = 4). Mice were deeply anesthetized with isoflurane (60-90 seconds) and then rapidly decapitated. Coronal slices containing the PFC or AM were cut in ice-cold solution containing (in mM) 92 NMDG, 20 HEPES, 25 Glucose, 30 NaHCO3, 1.2 NaH2PO4, 2.5 KCl, 5 Na-ascorbate, 3 Na-pyruvate, 2 Thiourea, 10 MgSO4, 0.5 CaCl2, saturated with 95% O2 5% CO2 (pH 7.3-7.4, ~305 mOsm/kg)] and incubated for 5-10 minutes at 35 degrees Celsius in the same solution. Slices were allowed to recover for a minimum of 30 minutes at room temperature in artificial cerebrospinal fluid (ACSF) containing (in mM) 126 NaCl, 2.5 KCl, 1.2 MgCl2, 2.4 CaCl2, 1.2 NaH2PO4, 21.4 NaHCO3, 11.1 Glucose, 3 Na-pyruvate, 1 Na-ascorbate. Recordings were made at 32-35°Celsius in the same solution which was bath perfused at 2-3 ml/min. For whole cell voltage clamp recordings, intracellular solution contained (in mM) 120 mM CsMeSO3, 5 mM NaCl, 10 mM TEA-Cl, 10 mM HEPES, 4 mM QX-314, 1.1 mM EGTA, 4 mM Mg-ATP, 0.3 mM Na-GTP (pH 7.2-7.3, ~290 mOsm/kg).

Whole cell recordings were performed in YFP-expressing PFC neurons projecting to AM, or in YFP-expressing AM neurons projecting to PFC. Differential interference contrast optics were used to patch neurons. For ChR2 experiments, a 473 nM laser (OEM laser systems, maximum output 500 mW) attached to fiber optic cable was used to deliver light to the slice. Light intensity of 8-12 mW was used to stimulate ChR2-expressing terminals in slice recordings. For experiments testing the involvement of glutamatergic transmission in the AM-projecting PFC neurons or in the PFC-projecting AM neurons, we measured optogenetically evoked EPSCs (Fig 6c-e) by giving 2 ms light pulses at a frequency of 0.5 Hz. In cells that showed a light-evoked response, DNQX (30 μM) and AP5 (50 μM) were added to the recording bath solution after a stable baseline measurement was recorded. Recordings were discarded if series resistance or input resistance changed >20% throughout the course of the recording. An Axopatch 200B amplifier (Molecular Devices) and Axograph X software (Axograph Scientific) were used to record and collect the data, which were filtered at 10 kHz and digitized at 4-20 kHz.

### Fiber-photometry calcium-signal recordings with optogenetic stimulation

We used the fiber photometry system manufactured by Doric Lenses (Quebec, Canada). The light pulse generator, consisting of a driver and LED units, produced light wavelengths (465 nm and 405 nm) in sinusoidal waveforms (208 Hz and 530 Hz, respectively), which were fed into a fluorescence mini-cube via patchcables (NA: 0.48; core diameter: 400 μm). The mini-cube combined these wavelength lights and sent the combined beam into the brain via a patchcable (NA: 0.48; core diameter: 400 μm) connected to the implanted optic fiber (NA: 0.48; core diameter: 200 or 400 μm). The same patchcable/optic fiber assembly, in turn, carried the emission of GCaMP (525 nm) as well as control (430 nm) back to the mini-cube, which then separated emission bandpass with beamsplitters and sent them to photoreceiver modules (Newport: model 2151) via patchcables (NA: 0.48; core diameter: 600 μm). The photoreceiver modules quantified signals and sent them to the Fiber Photometry Console, which was controlled by the Doric Neuroscience Studio software and which synchronized the acquisition of the data with the output of the laser stimulus. Recording data were collected at the rate of 1,200 Hz.

Med Associates’ system (Fairfax, VT) produced the trains (1, 2, 4, or 8 pulses at 25 Hz) delivered in a random order on a variable-interval schedule with the mean interval of 15 s by controlling the pulse generator (Doric Lenses) that, in turn, controlled a laser for generating a 3-ms pulse of a 635-nm wavelength. Each mouse received 41 trains for each pulse (41 trains x 4 train types =164 total trains) at the AM, VStr and VTA. Half of the mice received the trains at the AM first and then the VStr, while the other half received the trains in the reverse order. All mice received trains at the VTA after being tested at the AM and the VStr. For the delivery of VTA stimulation, the laser was connected to the mini-cube, which relayed lights to the VTA via the same probe for both the fiber photometry and optogenetic procedures. For mPFC stimulation, the laser was directly connected to the optic-fiber probe targeting the mPFC.

#### Photometry Analysis

We used custom-written MATLAB code to transform and analyze photometry data. We first binned the separate GCaMP and Control channels into one-minute epochs. We transformed the Control channels of each bin to a linear fit of its respective GCaMP channel, and calculated dF/F for each bin with the formula (GCaMP-FittedControl)/FittedControl^67^. We then extracted the dF/F trace corresponding to two seconds before and after every light train, extracted the corresponding area under the curve, and performed a repeated measures ANOVA with Time (before and after stimulation train) and Pulse (1, 2, 4, and 8 pulses) (GraphPad Prism) on the extracted area under the curve for each region of individual animal. We performed posthoc t-tests with Benjamini and Hochberg correction on Time for each pulse when the interaction was significant.

### Histology

Mice were intracardially perfused with 1X PBS followed by 4% paraformaldehyde. Brains were kept in a 30% sucrose in PBS solution over 1 – 2 days before cutting. After freezing, brains were coronally sectioned at 40 μm with a cryostat, mounted directly onto slides, and cover slipped with DAPI nuclear counterstain. Optical fiber placements and fluorophore expression were determined with fluorescent microscopy.

### Statistical analysis

Behavioral data were analyzed with Statistica or GraphPad Prism v4. Electrophysiological recording data were analyzed with Molecular Devices’ Clampfit v10.6 and Origin Pro v9.2 (OriginLab Corporation, MA, USA).

## Acknowledgments

The authors thanks NIDA Transgenic Breeding Facility for transgenic mice, NIDA Genetic Engineering and Viral Vector Core for viral vectors and NIDA Histology Core for cell counting.

## Author contributions

Conceptualization: S.I., Y.Y., Y.H.; Formal Analysis: Y.Y., Y.H., R.F.D. and S.I.; Investigation: C.Y., Y.H., A.D.T., C.T.P., L.A.R., A.J.K., R.F.D., S.J., A.T., A.F.P., C.N., C.B.C., C.S.; Software: R.F.D., C.M.-A.; Writing – Original Draft: S.I.; Writing - Review Editing: C.Y., Y.H., A.D.T., C.T.P., L.A.R., A.J.K., R.F.D., S.J., A.T., A.F.P., C.N., C.B.C., C.S., C.M.-A., D.V.W, H.L., Y.Y., and S.I.; Supervision, H.L., D.V.W., Y.Y. and S.I.; Projection Administration: S.I.

## Data and materials availability

All data is available in the main text or the supplementary materials.

## REFERENCES

1. Miller EK, Cohen JD. An integrative theory of prefrontal cortex function. Annu Rev Neurosci 24, 167–202 (2001).

2. Bechara A, Damasio AR, Damasio H, Anderson SW. Insensitivity to future consequences following damage to human prefrontal cortex. Cognition 50, 7–15 (1994).

3. Ridderinkhof KR, Ullsperger M, Crone EA, Nieuwenhuis S. The role of the medial frontal cortex in cognitive control. Science 306, 443–447 (2004).

4. Gilbert SJ, Burgess PW. Executive function. Current Biology 18, R110–R114 (2008).

5. Drevets WC, et al. Subgenual prefrontal cortex abnormalities in mood disorders. Nature 386, 824–827 (1997).

6. Mayberg HS. Limbic-cortical dysregulation: a proposed model of depression. J Neuropsychiatry Clin Neurosci 9, 471–481 (1997).

7. Kalivas PW, Volkow ND. New medications for drug addiction hiding in glutamatergic neuroplasticity. Mol Psychiatry 16, 974–986 (2011).

8. Everitt BJ, Robbins TW. Neural systems of reinforcement for drug addiction: from actions to habits to compulsion. Nat Neurosci 8, 1481–1489 (2005).

9. Mayberg HS, et al. Deep brain stimulation for treatment-resistant depression. Neuron 45, 651–660 (2005).

10. Downar J, et al. Anhedonia and reward-circuit connectivity distinguish nonresponders from responders to dorsomedial prefrontal repetitive transcranial magnetic stimulation in major depression. Biol Psychiatry 76, 176–185 (2014).

11. Terraneo A, Leggio L, Saladini M, Ermani M, Bonci A, Gallimberti L. Transcranial magnetic stimulation of dorsolateral prefrontal cortex reduces cocaine use: A pilot study. Eur Neuropsychopharmacol 26, 37–44 (2016).

12. Diana M, Raij T, Melis M, Nummenmaa A, Leggio L, Bonci A. Rehabilitating the addicted brain with transcranial magnetic stimulation. Nat Rev Neurosci 18, 685–693 (2017).

13. Dandekar MP, Fenoy AJ, Carvalho AF, Soares JC, Quevedo J. Deep brain stimulation for treatment-resistant depression: an integrative review of preclinical and clinical findings and translational implications. Mol Psychiatry 23, 1094–1112 (2018).

14. Menon V. Large-scale brain networks and psychopathology: a unifying triple network model. Trends Cogn Sci 15, 483–506 (2011).

15. Hebb DO. Drives and the C.N.S. (conceptual nervous system). Psychological review 62, 243–254 (1955).

16. Haber SN, Knutson B. The reward circuit: Linking primate anatomy and human imaging. Neuropsychopharmacology 35, 4–26 (2010).

17. Beier KT, et al. Circuit Architecture of VTA Dopamine Neurons Revealed by Systematic Input-Output Mapping. Cell 162, 622–634 (2015).

18. Ikemoto S, Yang C, Tan A. Basal ganglia circuit loops, dopamine and motivation: A review and enquiry. Behav Brain Res 290, 17–31 (2015).

19. Britt JP, Benaliouad F, McDevitt RA, Stuber GD, Wise RA, Bonci A. Synaptic and behavioral profile of multiple glutamatergic inputs to the nucleus accumbens. Neuron 76, 790–803 (2012).

20. Ikemoto S, Panksepp J. The role of nucleus accumbens dopamine in motivated behavior: a unifying interpretation with special reference to reward-seeking. Brain Res Rev 31, 6–41 (1999).

21. Taber MT, Fibiger HC. Electrical stimulation of the medial prefrontal cortex increases dopamine release in the striatum. Neuropsychopharmacology 9, 271–275 (1993).

22. Karreman M, Moghaddam B. The prefrontal cortex regulates the basal release of dopamine in the limbic striatum: an effect mediated by ventral tegmental area. J Neurochem 66, 589–598 (1996).

23. You ZB, Tzschentke TM, Brodin E, Wise RA. Electrical stimulation of the prefrontal cortex increases cholecystokinin, glutamate, and dopamine release in the nucleus accumbens: an in vivo microdialysis study in freely moving rats. J Neurosci 18, 6492–6500 (1998).

24. Russo SJ, Nestler EJ. The brain reward circuitry in mood disorders. Nat Rev Neurosci 14, 609–625 (2013).

25. Scofield MD, et al. The Nucleus Accumbens: Mechanisms of Addiction across Drug Classes Reflect the Importance of Glutamate Homeostasis. Pharmacol Rev 68, 816–871 (2016).

26. Alexander GE, Crutcher MD. Functional architecture of basal ganglia circuits: neural substrates of parallel processing. Trends Neurosci 13, 266–271 (1990).

27. Damasio AR. The somatic marker hypothesis and the possible functions of the prefrontal cortex. Philos Trans R Soc Lond B Biol Sci 351, 1413–1420 (1996).

28. Hare TA, Camerer CF, Rangel A. Self-control in decision-making involves modulation of the vmPFC valuation system. Science 324, 646–648 (2009).

29. Uylings HBM, Groenewegen HJ, Kolb B. Do rats have a prefrontal cortex? Behavioural Brain Research 146, 3–17 (2003).

30. Öngür D, Price JL. The organization of networks within the orbital and medial prefrontal cortex of rats, monkeys and humans. Cerebral Cortex 10, 206–219 (2000).

31. Alexander GE, DeLong MR, Strick PL. Parallel organization of functionally segregated circuits linking basal ganglia and cortex. Annu Rev Neurosci 9, 357–381 (1986).

32. Moorman DE, James MH, McGlinchey EM, Aston-Jones G. Differential roles of medial prefrontal subregions in the regulation of drug seeking. Brain Res 1628, 130–146 (2015).

33. Patton MH, Bizup BT, Grace AA. The infralimbic cortex bidirectionally modulates mesolimbic dopamine neuron activity via distinct neural pathways. J Neurosci 33, 16865–16873 (2013).

34. Vidal-Gonzalez I, Vidal-Gonzalez B, Rauch SL, Quirk GJ. Microstimulation reveals opposing influences of prelimbic and infralimbic cortex on the expression of conditioned fear. Learn Mem 13, 728–733 (2006).

35. Mayberg HS, et al. Regional metabolic effects of fluoxetine in major depression: serial changes and relationship to clinical response. Biol Psychiatry 48, 830–843 (2000).

36. Ikemoto S. Involvement of the olfactory tubercle in cocaine reward: intracranial self-administration studies. J Neurosci 23, 9305–9311 (2003).

37. Olds ME, Olds J. Approach-avoidance analysis of rat diencephalon. J Comp Neurol 120, 259–295 (1963).

38. Wright NF, Vann SD, Erichsen JT, O’Mara SM, Aggleton JP. Segregation of parallel inputs to the anteromedial and anteroventral thalamic nuclei of the rat. J Comp Neurol 521, 2966–2986 (2013).

39. Oh SW, et al. A mesoscale connectome of the mouse brain. Nature 508, 207–214 (2014).

40. van Groen T, Kadish I, Wyss JM. Efferent connections of the anteromedial nucleus of the thalamus of the rat. Brain Res Rev 30, 1–26 (1999).

41. Sun Q, et al. A whole-brain map of long-range inputs to GABAergic interneurons in the mouse medial prefrontal cortex. Nat Neurosci, (2019).

42. Murugan M, et al. Combined Social and Spatial Coding in a Descending Projection from the Prefrontal Cortex. Cell 171, 1663–1677 e1616 (2017).

43. McFarland K, Lapish CC, Kalivas PW. Prefrontal glutamate release into the core of the nucleus accumbens mediates cocaine-induced reinstatement of drug-seeking behavior. J Neurosci 23, 3531–3537 (2003).

44. Chen Q. The biggest trend in Chinese social media is dying, and another has already taken its place. Preprint at https://www.cnbc.com/2018/09/19/short-video-apps-like-douyin-tiktok-are-dominating-chinese-screens.html (2018).

45. Friston KJ, Buechel C, Fink GR, Morris J, Rolls E, Dolan RJ. Psychophysiological and modulatory interactions in neuroimaging. Neuroimage 6, 218–229 (1997).

46. Friston KJ, et al. Bayesian model reduction and empirical Bayes for group (DCM) studies. Neuroimage 128, 413–431 (2016).

47. Zeidman P, et al. A guide to group effective connectivity analysis, part 2: Second level analysis with PEB. Neuroimage 200, 12–25 (2019).

48. Ilango A, Kesner AJ, Broker CJ, Wang DV, Ikemoto S. Phasic excitation of ventral tegmental dopamine neurons potentiates the initiation of conditioned approach behavior: parametric and reinforcement-schedule analyses. Front Behav Neurosci 8, 155 (2014).

49. Ilango A, Kesner AJ, Keller KL, Stuber GD, Bonci A, Ikemoto S. Similar roles of substantia nigra and ventral tegmental dopamine neurons in reward and aversion. J Neurosci 34, 817–822 (2014).

50. de Bourbon-Teles J, et al. Thalamic control of human attention driven by memory and learning. Curr Biol 24, 993–999 (2014).

51. Wright NF, Vann SD, Aggleton JP, Nelson AJ. A critical role for the anterior thalamus in directing attention to task-relevant stimuli. J Neurosci 35, 5480–5488 (2015).

52. Lieberman MD, Straccia MA, Meyer ML, Du M, Tan KM. Social, self, (situational), and affective processes in medial prefrontal cortex (MPFC): Causal, multivariate, and reverse inference evidence. Neurosci Biobehav Rev 99, 311–328 (2019).

53. Gabbott PL, Warner TA, Jays PR, Salway P, Busby SJ. Prefrontal cortex in the rat: projections to subcortical autonomic, motor, and limbic centers. J Comp Neurol 492, 145–177 (2005).

54. Ikemoto S. Dopamine reward circuitry: two projection systems from the ventral midbrain to the nucleus accumbens-olfactory tubercle complex. Brain Res Rev 56, 27–78 (2007).

55. Ikemoto S. Brain reward circuitry beyond the mesolimbic dopamine system: a neurobiological theory. Neurosci Biobehav Rev 35, 129–150 (2010).

56. Akiguchi I, et al. Acute-onset amnestic syndrome with localized infarct on the dominant side--comparison between anteromedial thalamic lesion and posterior cerebral artery territory lesion. Jpn J Med 26, 15–20 (1987).

57. Ghika-Schmid F, Bogousslavsky J. The acute behavioral syndrome of anterior thalamic infarction: a prospective study of 12 cases. Ann Neurol 48, 220–227 (2000).

58. Shabbir SH, Nadeem F, Labovitz D. Anteromedial thalamic infarct: a rare presentation. BMJ Case Rep 2018, (2018).

59. Levy R, Dubois B. Apathy and the functional anatomy of the prefrontal cortex-basal ganglia circuits. Cereb Cortex 16, 916–928 (2006).

60. Xu M, Zhang SY, Dan Y, Poo MM. Representation of interval timing by temporally scalable firing patterns in rat prefrontal cortex. Proc Natl Acad Sci U S A 111, 480–485 (2014).

61. Wang DV, Yau HJ, Broker CJ, Tsou JH, Bonci A, Ikemoto S. Mesopontine median raphe regulates hippocampal ripple oscillation and memory consolidation. Nat Neurosci 18, 728–735 (2015).

62. Castagné V, Moser P, Roux S, Porsolt RD. Rodent models of depression: Forced swim and tail suspension behavioral despair tests in rats and mice. Current Protocols in Neuroscience, 1–14 (2011).

63. Lu H, Zou Q, Gu H, Raichle ME, Stein EA, Yang Y. Rat brains also have a default mode network. Proc Natl Acad Sci U S A 109, 3979–3984 (2012).

64. Power JD, Barnes KA, Snyder AZ, Schlaggar BL, Petersen SE. Spurious but systematic correlations in functional connectivity MRI networks arise from subject motion. Neuroimage 59, 2142–2154 (2012).

65. Mai JK, Majtanik M, Paxinos G. Atlas of the human brain, 4th edn. Academic Press (2015).

66. Friston KJ, Harrison L, Penny W. Dynamic causal modelling. Neuroimage 19, 1273–1302 (2003).

67. Calipari ES, et al. In vivo imaging identifies temporal signature of D1 and D2 medium spiny neurons in cocaine reward. Proc Natl Acad Sci U S A 113, 2726–2731 (2016).

68. Franklin KBJ, Paxinos G. The Mouse Brain in Stereotaxic Coordinates, 3rd edn. Elsevier (2007).

