## Supplementary Figures for "Medial Prefrontal Cortex Interacts with the Anteromedial Thalamus in Motivation and Dopaminergic Activity"

**This PDF file includes:**

Supplementary Figs. 1 to 9

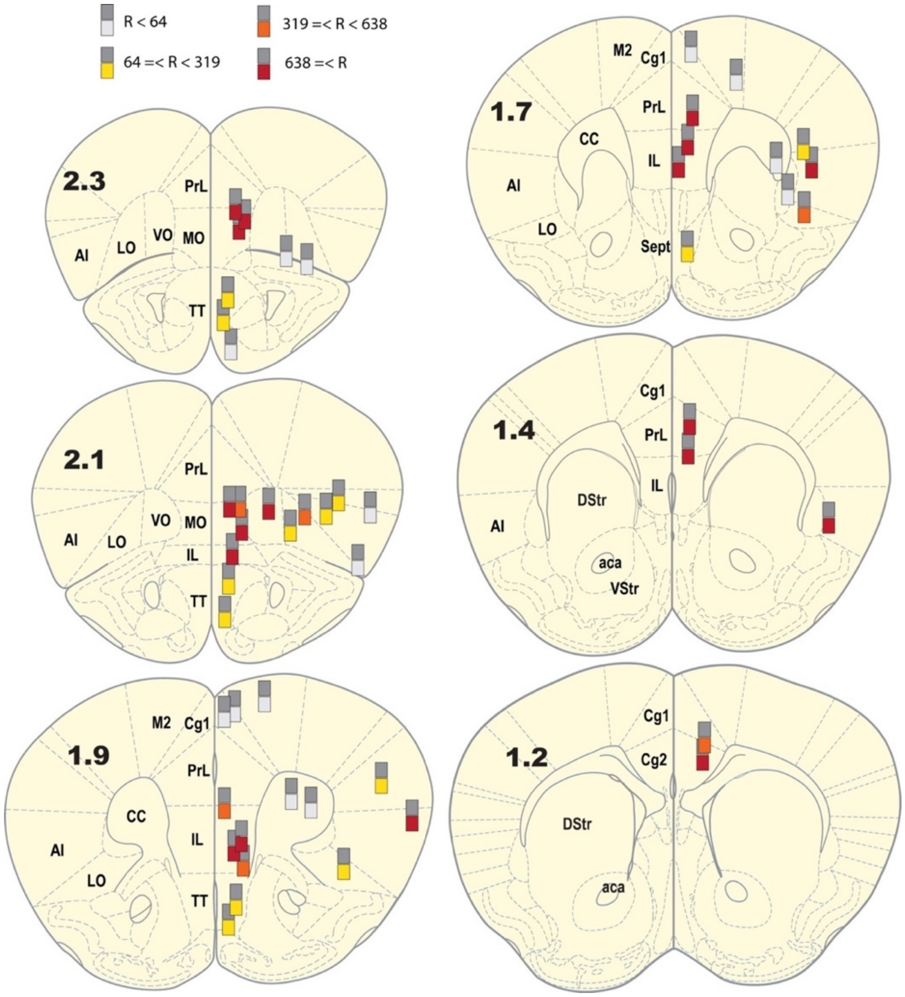


**Suppl. Fig. 1. The sites and effectiveness of optogenetic stimulation delivered at the PFC and TT in ICSS.** Each dark-gray square indicates the tip of optic fiber, which is accompanied with a colored square below, indicating one of four levels of ICSS rates. Coronal sections were adopted from the mouse brain atlas[1]. Abbreviations: AI, anterior insular cortex; CC, corpus collosum; Cg1, dorsal anterior cingulate cortex; Cg2, ventral anterior cingulate cortex; IL, infralimbic cortex; LO, lateral orbital cortex; M2, secondary motor cortex; MO, medial orbital cortex; PrL, prelimbic cortex; TT, tenia tecta; VO, ventral orbital cortex. Note that although some rodent brain atlases distinguish the IL from the dorsal peduncular cortex[1, 2], others do not[3, 4, 5], and the present study adopted the latter. Similarly, the AI on the drawing included the anterior insular, dorsal insular, granular insular, and dysgranular insular cortices.

References

1. Paxinos G, Watson C. *The Rat Brain in Stereotaxic Coordinates*, 6th edn (2007).
2. Franklin KBJ, Paxinos G. *The Mouse Brain in Stereotaxic Coordinates*, 3rd edn. Elsevier (2007).
3. Swanson LW. *Brain Maps: Structure of the Rat Brain* (1992).
4. Uylings HBM, Groenewegen HJ, Kolb B. Do rats have a prefrontal cortex? *Behavioural Brain Research* **146**, 3-17 (2003).
5. Öngür D, Price JL. The organization of networks within the orbital and medial prefrontal cortex of rats, monkeys and humans. *Cerebral Cortex* **10**, 206-219 (2000).

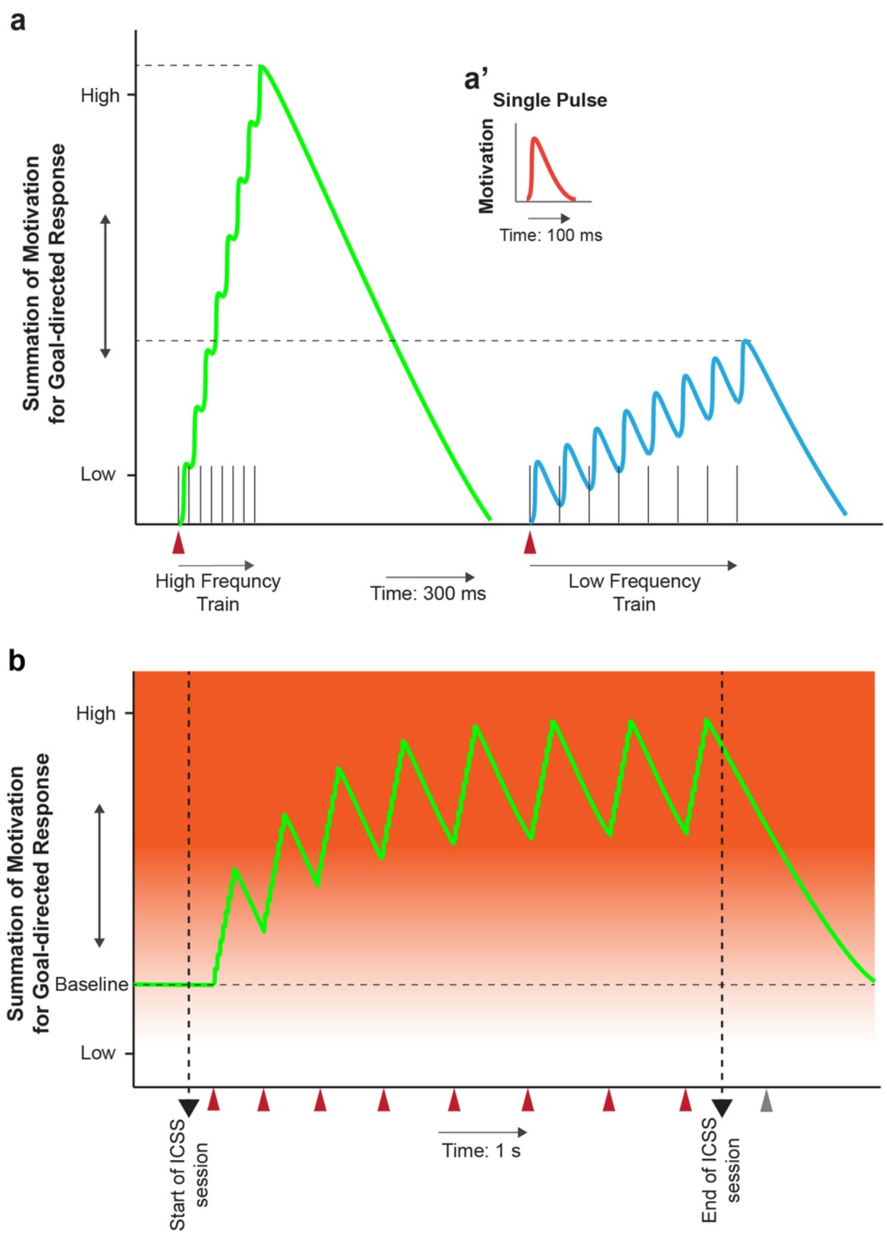


**Suppl. Fig. 2. Theoretical models explaining how pulse interval and train interval play important roles in ICSS rates and goal-directed motivation. a** Model depicting the summation effects on motivation of a high-frequency and a low-frequency trains of 8 pulses. Each pulse triggers a small motivation effect (a'), which decays quickly. When pulses are delivered close in time, they summate to elicit a significant effect on motivation. The arrowhead indicates the onset of a pulse train. **b** Model depicting the summation effect of trains delivered closely in time. When trains occur close in time, they increase motivation for goal-directed behavior, resulting in a high rate of lever responding. The arrowheads with maroon and gray colors indicate the time of lever response reinforced and not reinforced, respectively. Similar models are suggested to explain the relationship between dopamine and ICSS rate[6].

6. Ikemoto S, Yang C, Tan A. Basal ganglia circuit loops, dopamine and motivation: A review and enquiry. *Behav Brain Res* **290**, 17-31 (2015).

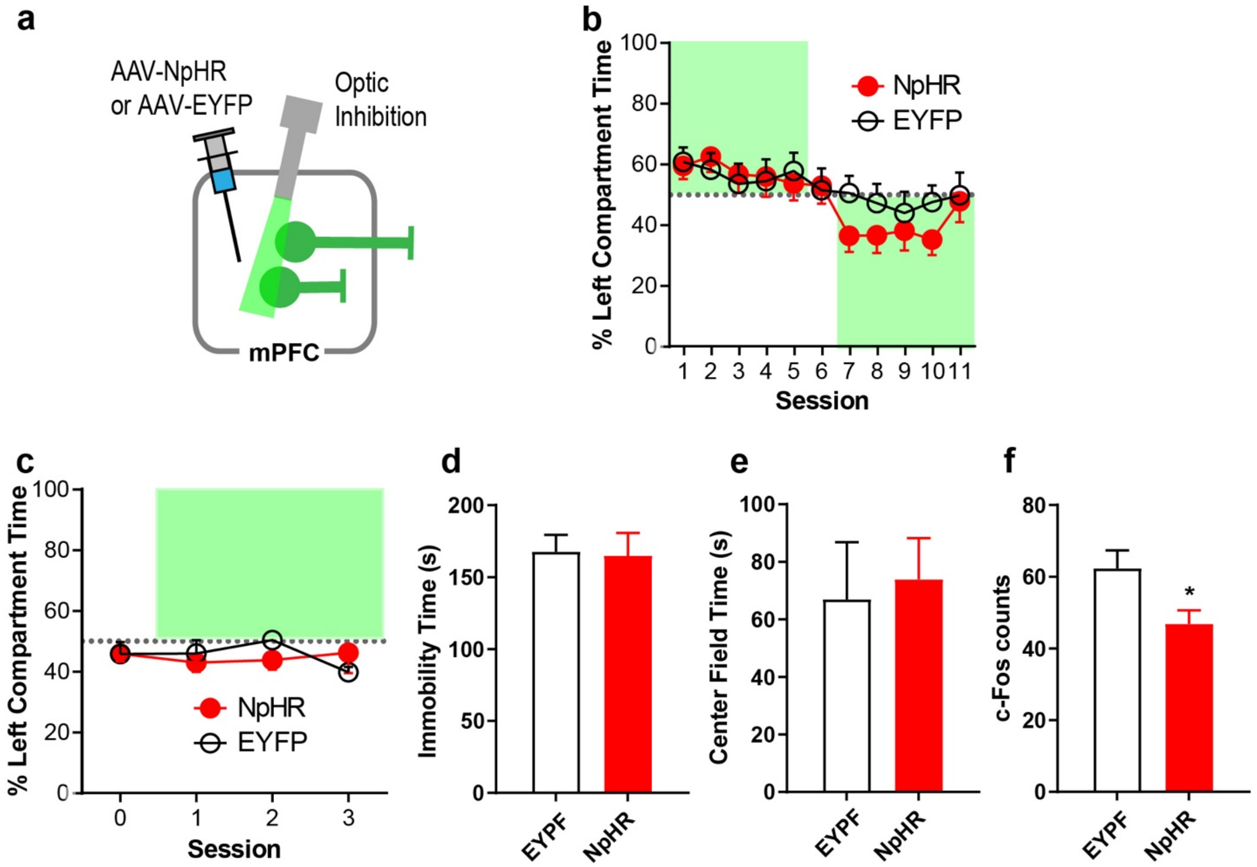


**Suppl. Fig. 3. Effects of the inhibition of mPFC neurons on affective state.** The results described in this figure suggest that tonic activity of mPFC neurons is not involved in suppressing negative emotional states. **a** Mice received an injection of AAV encoding the gene for halorhodopsin-3.0 (NpHR) or EYFP only and an optic fiber at the mPFC. **b** Real-time place-preference data with unilateral inhibition. NpHR: n = 6; EYFP: n = 6. Data (means ± SEM) are derived by the equation: (left compartment time – right time) / (left time + right time) * 100. Data points fell in green squares indicate relative preference to the stimulation compartment. **c-f** Bilateral inhibition data with two groups NpHR (n = 5) and EYFP (n =4). **c** Real-time place-preference data. Photostimulation was delivered in the left compartment for sessions 1-3, but not session 0. Data (means ± SEM) are derived by the equation: (left compartment time – right time) / (left time + right time) * 100. **d** Mean immobility times (± SEM) during forced swim. **e** Mean center-field times (± SEM) in open field. **f** To confirm that the NpHR manipulation inhibited mPFC neurons, the mice were placed in a novel chamber for 30 min with mPFC photostimulation before being euthanized. mPFC c-Fos expressions were then compared between the experimental and control groups. The data are mean c-Fos counts (± SEM) in the mPFC. The NpHR group had significantly less c-Fos expression than the control group (*$P <$ 0.05), suggesting that the photostimulation inhibited mPFC neurons during the tests described above.

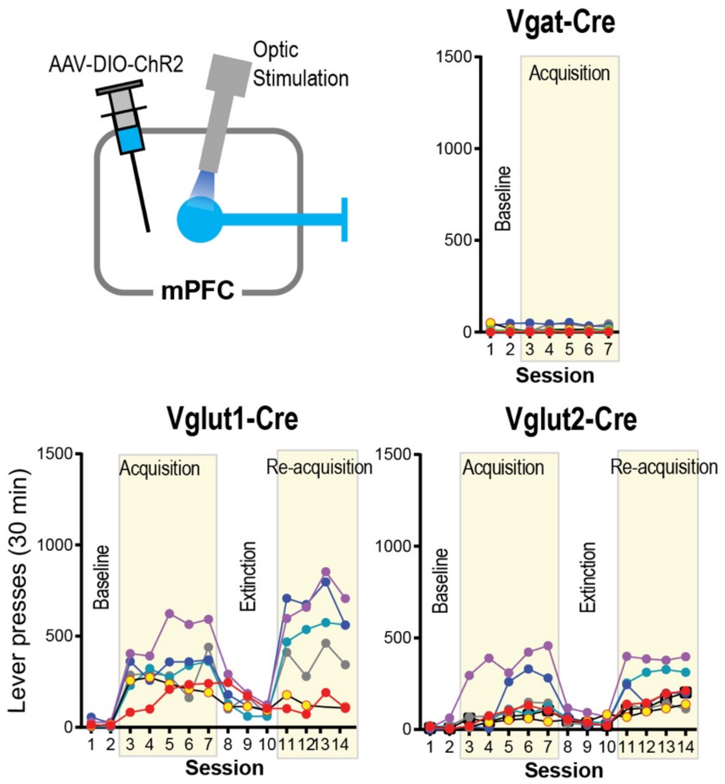


**Suppl. Fig. 4. Glutamatergic neurons are important in goal-directed motivation induced by mPFC stimulation.** While most cortical glutamate neurons are vesicular glutamate transporter type 1 (Vglut1), the vicinity of the IL contains Vglut2 neurons. Cre-dependent AAV-ChR2-EYFP was injected into the mPFC followed by the implantation of an optic fiber at the same region of transgenic mice: Vlgut1-Cre, Vglut2-Cre, or Vgat-Cre mice. The mice received photostimulation (a train of 8 pulses at 25 Hz) in sessions 3-7 and 11-14, while no photostimulation in sessions 1-2 or 8-10. mPFC photostimulation reinforced responding of all Vglut1-Cre (n = 6) and Vglut2-Cre (n = 6) mice, whereas the same stimulation failed to reinforce responding of Vgat-Cre mice (n = 6).

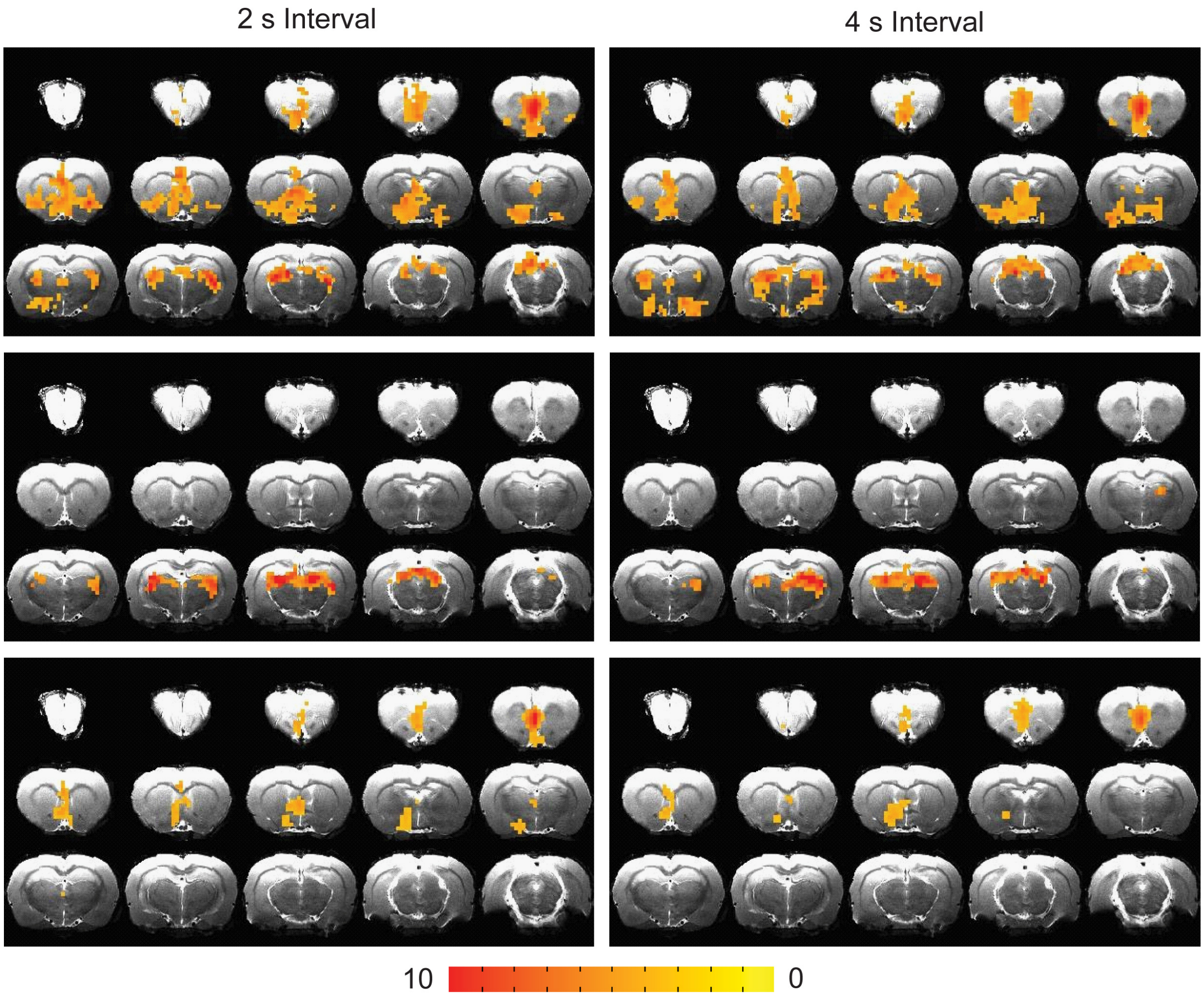


**Suppl. Fig. 5. fMRI: mPFC stimulation with 2 or 4-s interval increases less BOLD signals throughout the brain.** The photostimulation with the 2- or 4-s interval increased BOLD signals in extensive subcortical regions of the mPFC with ChR2 (top; n = 10), while BOLD signals in the control group were confined in the visual thalamus and superior colliculus (middle; n = 7). The 2- or 4-s interval stimulation resulted in significant differences in BOLD signal between the ChR2 and control groups in fewer regions (bottom) than the 1-s interval stimulation (Fig. 3e, bottom).

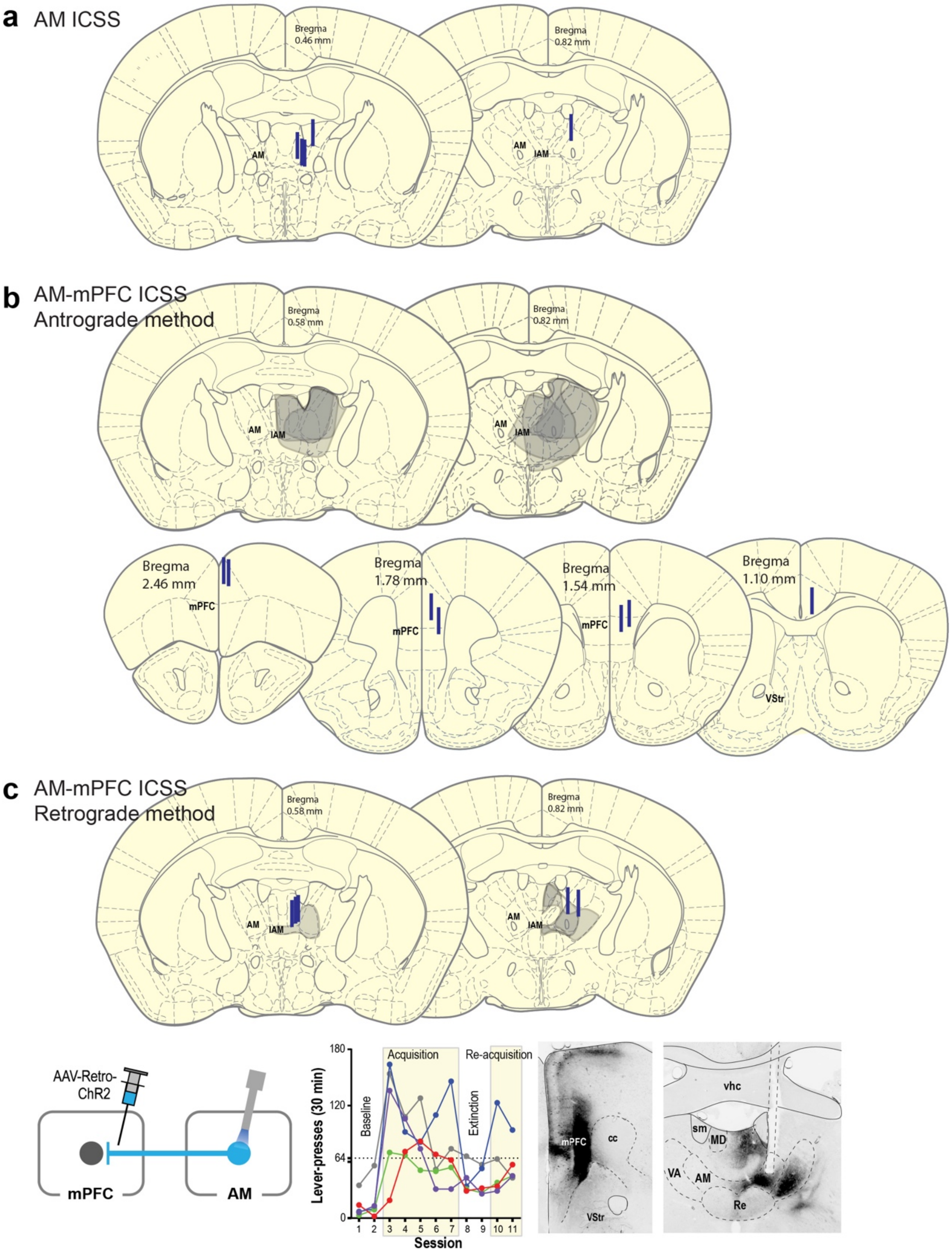


**Suppl. Fig. 6. Sites showing AAV injections and optic-fiber implantations for AM experiments.** Drawings of coronal sections showing optic-fiber placements for AM stimulation (**a**), areas affected by AAV injections (**b**, top) and optic-fiber placements for mPFC stimulation (**b**, bottom), and areas affected by AAV injections and optic-fiber placements for AM stimulation (**c**).

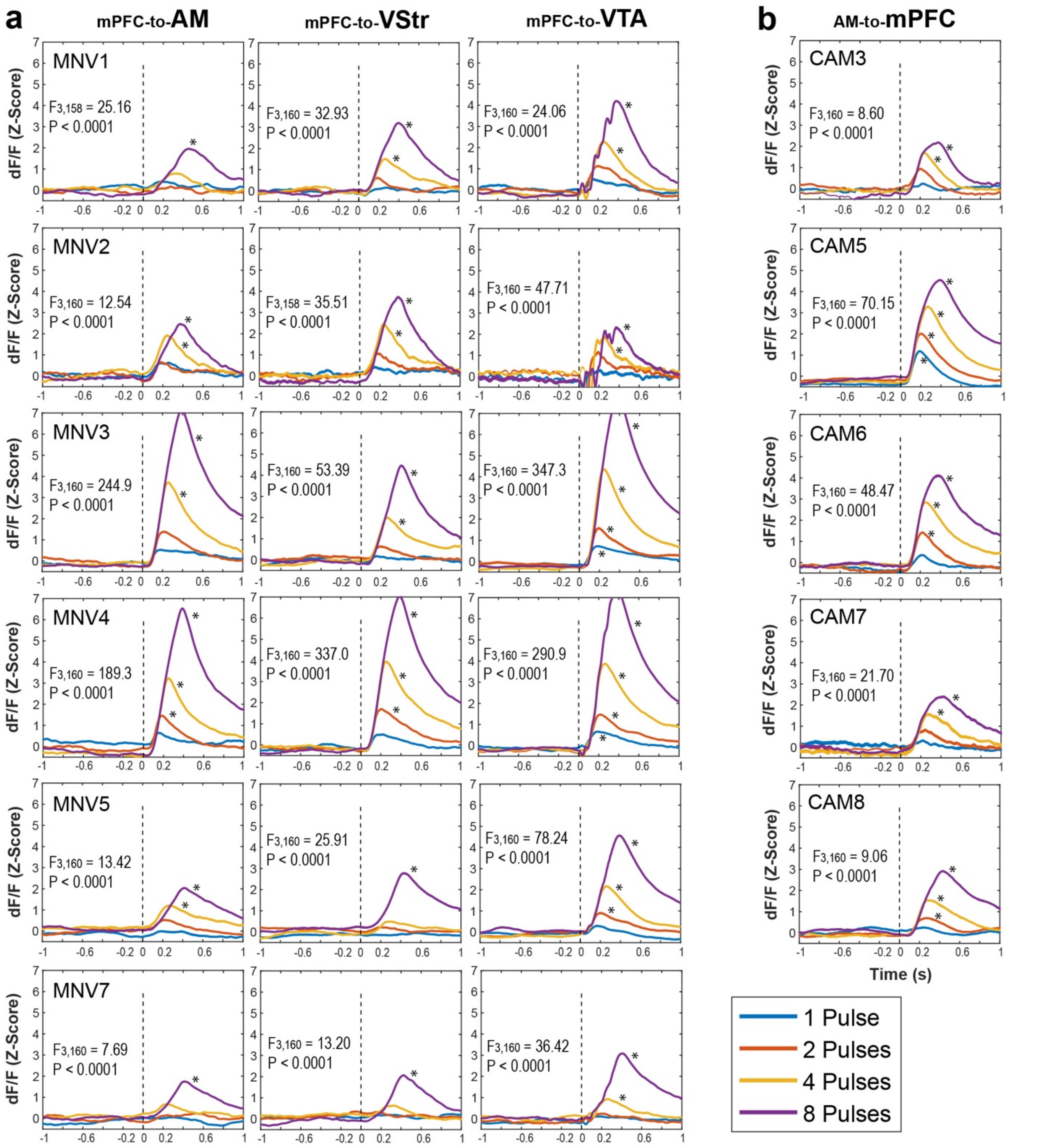


**Suppl. Fig. 7. Data of individual mouse data showing that the stimulation of mPFC axonal terminals at the AM or of AM axonal terminals at the mPFC activates VTA DA neurons.** GCaMP signals (Z-score dF/F) of VTA DA neurons induced by 4-pulse trains (1, 2, 4 and 8) delivered at the mPFC-to-AM, mPFC-to-VStr and mPFC-to-VTA pathways (**a**) and at the AM-to-mPFC pathway (**b**). The F- and P-values shown within each graph indicate the interaction with a 2-by-4 within-subjects ANOVA with Time (before and after stimulation train) and Pulse (1, 2, 4, and 8 pulses) on areas under curve for each region of each mouse. When the interaction was significant, posthoc t-tests with Benjamini and Hochberg correction were conducted on Time for each pulse. $^*P < 0.001$.

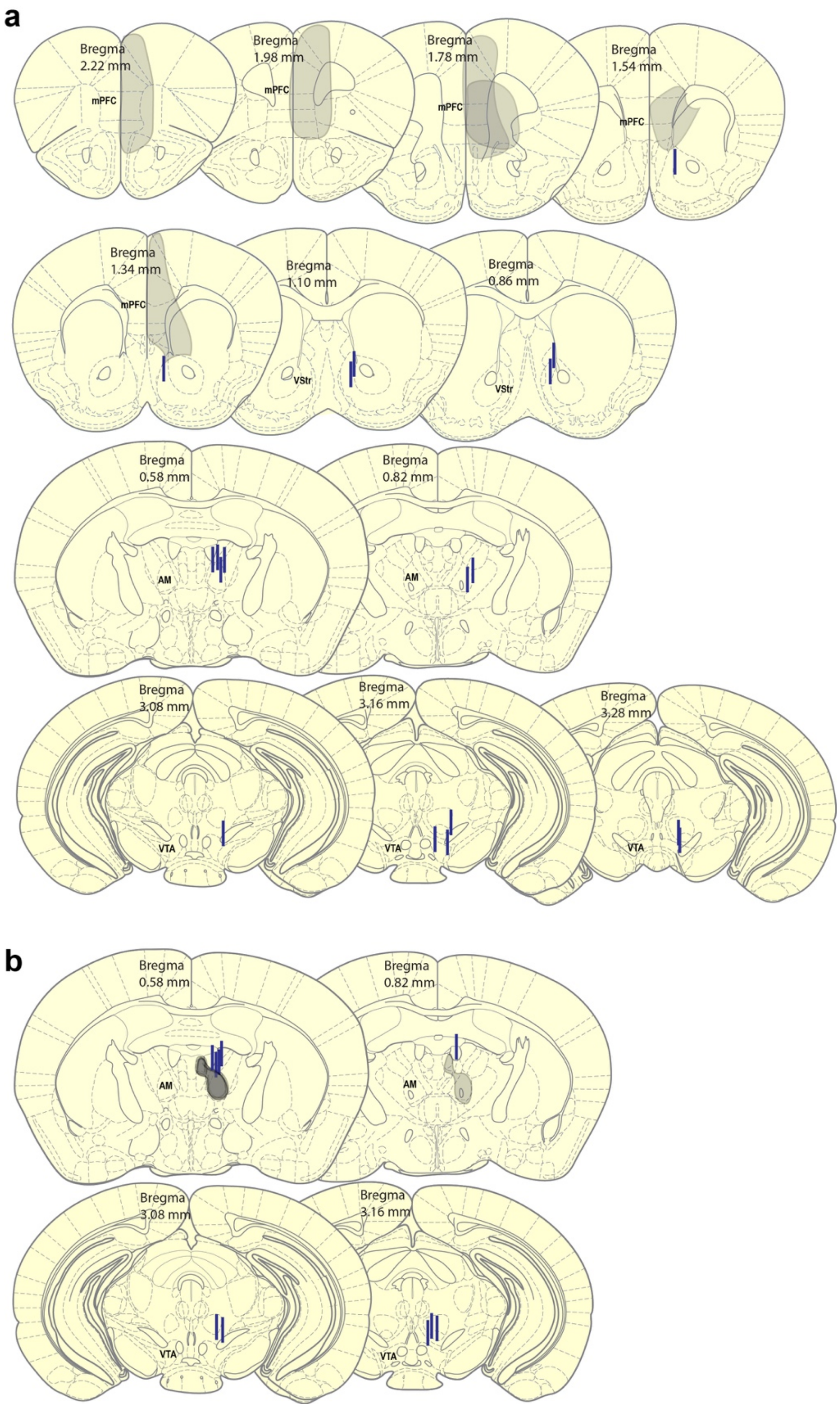


**Suppl. Fig. 8. Sites showing the expression of Chrimson and stimulation and recording sites.** Drawings of coronal sections showing areas affected by AAV injections and optic-fiber placements for the stimulation or recording of the VStr, AM, and VTA. **a** Results of the mice used for the stimulation of mPFC neuron terminals at the AM, VStr and VTA. **b** Results of the mice used for AM-to-mPFC stimulation.

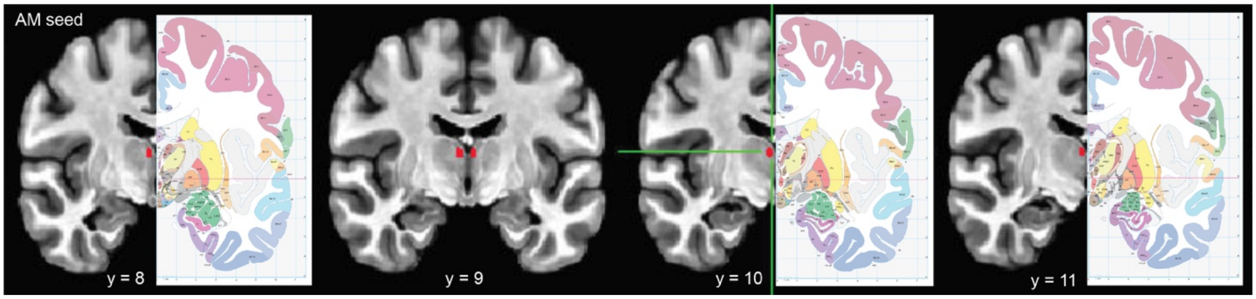


**Suppl. Fig. 9. AM seed**. Exact location of the AM (red circles) was determined and hand-drawn on the MNI_T1_152_2009_template with reference to *Atlas of The Human Brain*[7].

7. Mai JK, Majtanik M, Paxinos G. *Atlas of the human brain*, 4th edn. Academic Press (2015).
